# Mutually inhibiting teams of nodes: A predictive framework for structure–dynamics relationships in gene regulatory networks

**DOI:** 10.1101/2025.02.07.636066

**Authors:** Sai Shyam, Nikhil Nandhan, Vaibhav Anand, Mohit Kumar Jolly, Kishore Hari

## Abstract

Phenotypic plasticity—the reversible switching of cell-states—is a central tenet of development, regeneration, and cancer progression. These transitions are governed by gene regulatory networks (GRNs), whose topological features strongly influence their dynamics. While toggle switches (mutually inhibitory feedback loops between two transcription factors) are a common motif observed for binary cell-fate decisions, GRNs across diverse contexts often exhibit a more general structure: two mutually inhibiting teams of nodes. Here, we investigate the teams of nodes as a potential topological design principle of GRNs. We first analyze GRNs from the Cell Collective database and introduce a metric, impurity, which quantifies the fraction of edges inconsistent with an idealized two-team architecture. Impurity correlates strongly with statistical properties of GRN phenotypic landscapes, highlighting its predictive value. To further probe this relationship, we simulate artificial two-team networks (TTNs) using both continuous (RACIPE) and discrete (Boolean) formalisms across varying impurity, density, and network size values. TTNs exhibit toggle-switch-like robustness under perturbations and enable accurate prediction of dynamical features such as inter-team correlations and steady-state entropy. Together, our findings establish the teams paradigm as a unifying principle linking GRN topology to dynamics, with broad implications for inferring coarse-grained network properties from high-throughput sequencing data.

## Introduction

Phenotypic plasticity, the ability of cells to change their phenotypes reversibly in response to changing environments, is a fundamental phenomenon (Mills et al. 2019) across different contexts, including embryonic development and disease (Jopling et al. 2011; Gupta et al. 2019; Kalra et al. 2021; Pérez-González et al. 2023). Cell-state transitions often require a large-scale change in the transcriptomic profile, which are governed by the underlying gene regulatory networks (GRNs). Some topological features of GRNs, such as feedback and feedforward loops, have a significant impact on the dynamics of phenotypic plasticity(Murphy et al. 2002; Zhang et al. 2016; Hebbar et al. 2022). Many embryonic cell-fate transitions are governed by relative expression levels of two mutually repressing genes (Zhou and Huang 2011). This regulatory motif of two genes (A and B) mutually inhibiting each other is called a Toggle Switch (TS) and can give rise to two co-existing stable steady states that can switch back and forth: (high A, low B) and (low A, high B) (Gardner et al. 2000). For example, the master regulator genes PU.1 and GATA1 form a toggle switch that influences the differentiation of common myeloid progenitor cells into myeloid and erythroid progenitors based on the relative expression of these two genes. (High PU.1, low GATA1) facilitates the differentiation of the common myeloid progenitor cell into myeloid cells, while (low PU.1, high GATA1) leads to differentiation into erythroid cells. A third state, enabled by self-activation on the two transcription factors, has a balanced expression level of both PU.1 and GATA1 molecules and corresponds to the progenitor state (Roeder and Glauche 2006). Similar binary cell-fate decision systems are also important in cancer (Pérez-González et al., 2023; Sahoo et al., 2021). A prime example is epithelial-to-mesenchymal transition (EMT), where a toggle switch between the family of transcription factors ZEB1/2 and the family of miR-200 microRNAs has been identified to be a core motif in the underlying GRNs (Brabletz and Brabletz 2010; Celià-Terrassa et al. 2018). The self-activation of ZEB1 supports the emergence of a hybrid E/M state, composed of expression of both ZEB1/2 and miR200 molecules, similar to the progenitor state in the (PU.1, GATA1) example (Lu et al. 2013; Jolly et al. 2018). These hybrid E/M states are more stem-like, drug-resistant, and metastatically fit (Pastushenko and Blanpain 2019).

The dynamics of a toggle switch, including transcription factors and/or microRNAs, have been well-studied using deterministic and stochastic models to capture qualitative features of binary cell-fate decisions (Tian and Burrage 2006; Roeder and Glauche 2006; Shiraishi et al. 2010; Jaruszewicz and Lipniacki 2013; Okawa et al. 2016). Furthermore, the emergent properties of the toggle switch are resilient to external perturbations, making the toggle switch a robust cell-fate decision module (Harlapur et al. 2022). However, across different contexts, the underlying GRNs of cell-fate decisions have a larger number of nodes, encompassing multiple toggle switches and feedback loops (Ríos et al. 2015; Udyavar et al. 2017; Hari et al. 2020; Arthur et al. 2024). Recently, we analysed the networks governing EMT and found the presence of two mutually inhibiting teams of nodes (Hari et al. 2022b). A team is a group of genes that activate member genes belonging to the same team while inhibiting genes of other teams directly and/or indirectly. For instance, in the EMT network, most members of the epithelial team were EMT-inhibiting microRNAs such as miR-200 family, while most members of the mesenchymal team were EMT-inducing transcription factors (EMT-TFs) such as ZEB1 and SNAI1. These EMT-TFs activated each other and also repressed microRNAs that targeted them or other EMT-TFs, while many microRNAs inhibited EMT-TFs that inhibited them or other microRNAs in the same team. Such ‘teams’ have also been observed in other biological contexts, including developmental decision-making (Chauhan et al. 2021; Pillai and Jolly 2021; Sehgal et al. 2024; Hari et al. 2025). The intuitive framework of mutually inhibiting teams, their consistent influence on network dynamics, and their presence across diverse biological contexts make them an attractive candidate for being a new topological design principle of gene regulatory networks (GRNs). However, the role of network parameters and external perturbations in shaping the dynamics of team-based GRNs remains poorly understood. In particular, their impact on the phenotypic repertoire of decision-making GRNs has yet to be investigated comprehensively.

Here, we assess the potential of teams of nodes as a broadly applicable design principle by examining their occurrence in biological networks and probing how perturbations affect their emergent dynamics. We first analyzed GRNs from the Cell Collective database, which spans a wide range of biological systems and thus exhibited a wide variation in team strength values. To systematically characterize this variability, we introduced a metric called impurity, defined as the fraction of edges inconsistent with an idealized two-team architecture. Strikingly, impurity correlated with statistical properties of the phenotypic landscapes of Cell Collective GRNs. Building on this insight, we further investigated how impurity, network density, and size shape the dynamics of artificially generated team-based networks. We simulated artificially generated two-teamed networks (TTNs) using two simulation formalisms: primarily a threshold-based Boolean formalism, a discrete state-space approach that considers only the effect of the network topology without any parameter sets (Font-Clos et al. 2018), and RACIPE, a continuous state-space, ODE-based simulation formalism that calculates network dynamics over a large parameter space (Huang et al. 2018). We quantified two key properties of the phenotypic landscape to evaluate how network parameters shape emergent dynamics: (a) the frequency of pure states (F1)— mutually exclusive states in which one team is “on” and the other “off”—and (b) the correlation of expression levels between nodes belonging to opposing teams (CC), using the toggle switch as a reference. By mapping changes in these properties as a function of network parameters, we found consistent trends between artificial networks and those from the Cell Collective database. Our results further demonstrate that two-team networks (TTNs), much like toggle switches, preserve their dynamical behavior under external perturbations. Using artificial TTNs, we showed that constraining GRNs into a two-team structure enables accurate prediction of experimentally relevant dynamical features, such as node–group correlations and the entropy of steady-state distributions. Together, these findings establish the teams’ paradigm as a powerful framework for linking GRN structure to dynamics. Beyond predictive insights, this paradigm also offers a foundation for developing new methods to infer coarse-grained network properties from high-throughput data, advancing our understanding of emergent cellular behaviors.

## Results

### Teams of nodes and binary landscapes in cell-fate decision networks

In this section, we start with a brief introduction to the idea of teams, discussing the network properties, and the properties of the emergent landscapes with examples of cell fate decision networks that we had analyzed previously. Consider the cell-fate decision system – Epithelial-Mesenchymal plasticity (EMP). EMP is a crucial process studied in developmental biology and cancer biology, involving cell sampling phenotypes on a spectrum with epithelial phenotype (E) on one end and mesenchymal phenotype (M) on the other. The phenotypic landscape of EMP, typically representing the abundance of different phenotypes along the EMP spectrum in a population of heterogeneous cells, could be classified as predominantly binary. A salient feature of experimental studies involving EMP is that a heterogeneous cell population primarily would consist of cells with either E or M phenotype, with a small percentage corresponding to hybrid E/M phenotypes. Thus, the phenotypic frequency distribution predominantly looks bimodal. This feature of bimodality is observed across a wide variety of cell-fate decision systems, a few examples being NE/non-NE decision system in small cell lung cancer (SCLC), gonadal cell-fate determination system and a majority of decision systems involved in development. As described in the discussion earlier, such systems could be visualized as being driven by toggle switches, formed by two master regulators mutually inhibiting each other.

However, we found that the real networks are not as clean-cut. While some networks present high-degree nodes that could be viewed as the master regulators, other networks do not. In some cases, the markers of a given phenotype do not regulate each other at all, but are co-expressed due to the regulation by the markers of the other phenotype. Despite the individual differences in structure and hierarchy of these networks, we find that they can be united together as networks with teams, by considering not only the direct interactions, but also the indirect interactions. Consider the example in **Fig. 1A, i**), an EMP network which has been constructed to explain the phenotypic heterogeneity in a population of cells undergoing EMP. The nodes in the network have been segregated into two groups based on their biological functions: Epithelial and Mesenchymal. Notice how there are no direct interactions between the epithelial nodes (micro-RNAs, labelled as miRs). However, one can realize that the epithelial nodes are inhibiting mesenchymal nodes, and the mesenchymal nodes are, in turn, inhibiting epithelial nodes. Therefore, epithelial nodes are activating themselves indirectly. Inspired by this observation and the fact that the dynamics of a node carry the effect of the regulators of its regulators by proxy, we transform the structure of the network to better represent the dynamics. We integrate the effects of interactions of different path lengths between each pair of nodes, where a path is a series of connected edges between any pair of nodes, and the length of such a path is n-1, where n is the number of nodes in the path. Thus, we can say that while there are no paths of length 1 between Epithelial nodes, there are paths of length 2 mediated by mesenchymal nodes. Following this transformation, we can construct the “influence matrix”, which is also a signed, directed graph as the original network, but more connected. This influence matrix can be further clustered to reveal the “team” structure of a network, as shown in **Fig. 1A, ii, S1**). The team structure emerges from the network such that the nodes belonging to the same team have a positive influence on each other, while those belonging to the opposite team have a negative influence on each other. It is interesting to note that the influence matrix looks similar to that of a correlation or co-expression matrix, given its connectivity, despite being derived from the underlying regulatory network. Indeed, we were able to show previously that the influence matrix closely resembles the pairwise correlation matrix obtained from simulated steady states of the network.

**Figure 1:**
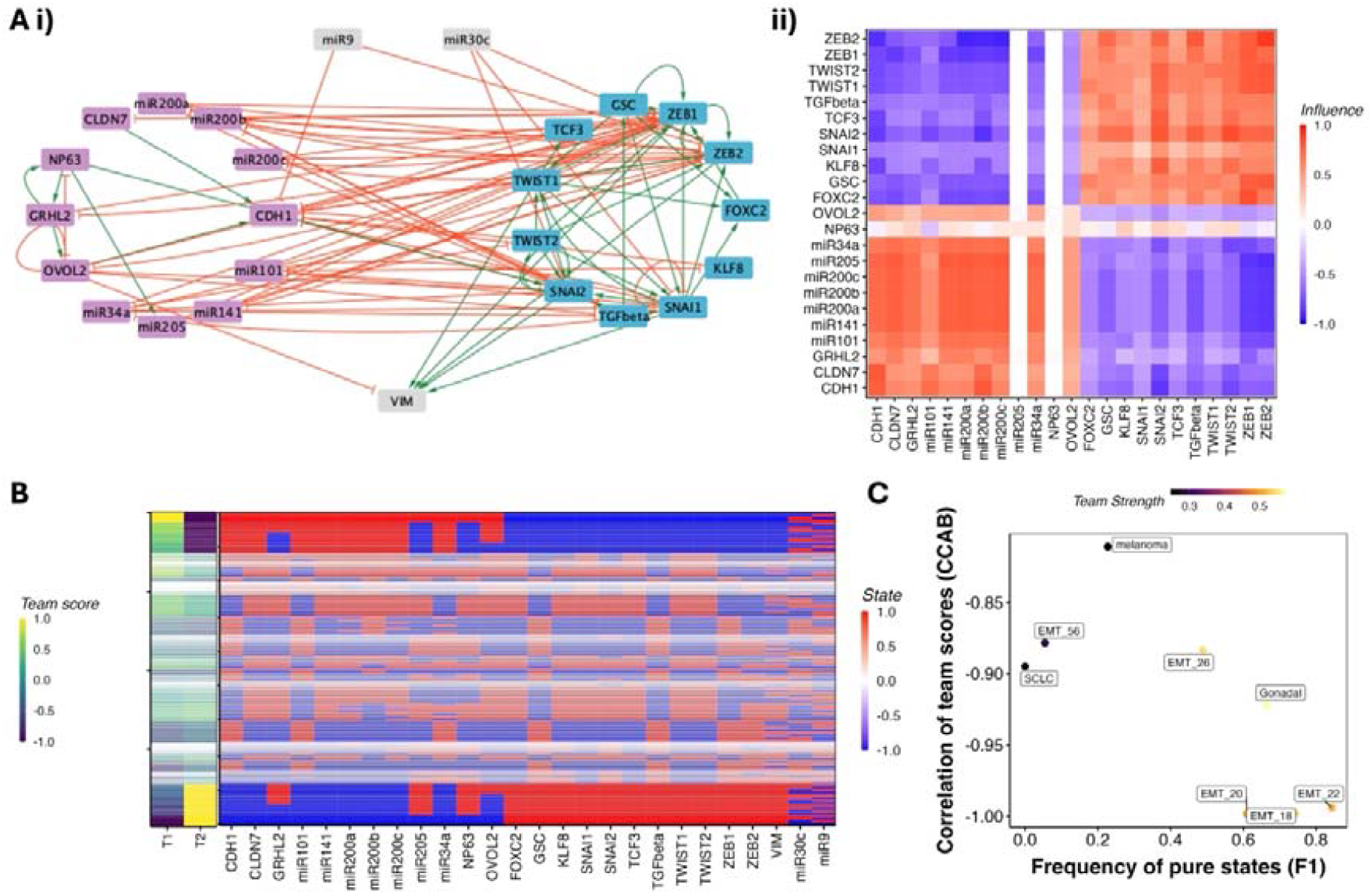
Teams and the phenotypic landscape of cell-fate decision networks: **A)** Network wiring diagram (i) and the influence matrix (ii) of a 26-node EMT network (Tripathi et al. 2020). **B)** Concatenated heatmaps depicting the steady state expression levels of the nodes in the EMT network (right) and corresponding team scores of each steady state (left). Each row corresponds to a steady state, and each column corresponds either to a node of the network or a team. The width of each row is proportional to log(frequency). **C)** F1 and CCAB of 8 cell fate decision networks that we have analyzed in (Hari et al. 2022b).

We simulated these networks using a threshold-based Boolean formalism (see methods) to obtain an ensemble of steady states that represents the experimental heterogeneous population. The influence matrix thus reveals an underlying toggle-switch with self-activation (TSSA) structure in the network, with each team acting as a single node in TSSA. Importantly, as one would expect from the TSSA, the majority of the state space of the converges to one of the two states where nodes from one team have a high expression and the other team have a low expression (**Fig. 1B**, red-blue matrix – thickness of the row shows the abundance of the steady state) – a binary phenotypic landscape. Using a metric that quantifies the team strength by measuring the strength of influences within and across teams (thereby measuring how well the network allows for a team to act as a single unit), we were able to qualitatively demonstrate that strong teams lead to a strongly bimodal phenotypic landscape. Using the analogy of teams acting as supernodes, we calculated the frequency of pure states (F1, representing the frequency of states with only one team on), which represents the total abundance of the two dominant phenotypes at the ends of the spectrum, and the correlation between the expression levels of the “supernode” (CCAB) calculated for each steady state as the average expression of all nodes in a given team. Networks with higher team strengths showed higher F1 scores and stronger correlations, and vice versa (**Fig. 1C**). Motivated by these observations, we wanted to ask if we can quantify the effect of network structure on the phenotypic landscape emergent from GRNs.

### Network impurity correlates with F1 and CCAB in GRNs from Cell Collective database

To test whether the existence of teams is a general feature of GRNs beyond the previously studied cases, we analyzed GRNs from the Cell Collective database (Helikar et al. 2012), spanning a range of biological contexts. We eliminated the networks having more than 100 nodes from this analysis for ease of computational tractability, leaving us with 42 networks in total. Of these, approximately 20 networks exhibited mutually inhibiting team structures of varying strengths (see **Fig. 2A, S2A**). In addition, we identified more complex interaction motifs between teams in some networks. For example, some networks exhibited mutually *activating* teams (e.g., signalling network that include links to propagate and amplify signals through positive feedback loops (der Heyde et al. 2014)), while others included negative feedback loops between teams (e.g., cell cycle networks capturing oscillations (Todd and Helikar 2012)). Given the similarity between the influence matrix and the corresponding correlation matrices observed previously, identification of these team-based interaction structures can offer an intuitive lens through which to understand the emergent phenotypic landscape of GRNs. Mutually inhibitory teams tend to result in two distinct, “pure” phenotypic states as shown above: one in which the nodes of one team are predominantly active (expressed), and another in which the opposite is true. On the other hand, mutually activating teams—or, equivalently, a single coherent team—tend to give rise to phenotypes characterized by the co-expression of all team members. The negative feedback was observed in a cell cycle network, where oscillations between cell cycle phases is a desired outcome and can be supported by negative feedback loops. However, we also observed that nearly half of the networks from the Cell Collective database did not exhibit clearly defined teams, and even among those that did, the team structures varied considerably in strength and organization. This led us to ask whether it might still be possible to represent these more ambiguous networks using a relaxed concept of “*impure* teams.”

**Figure 2:**
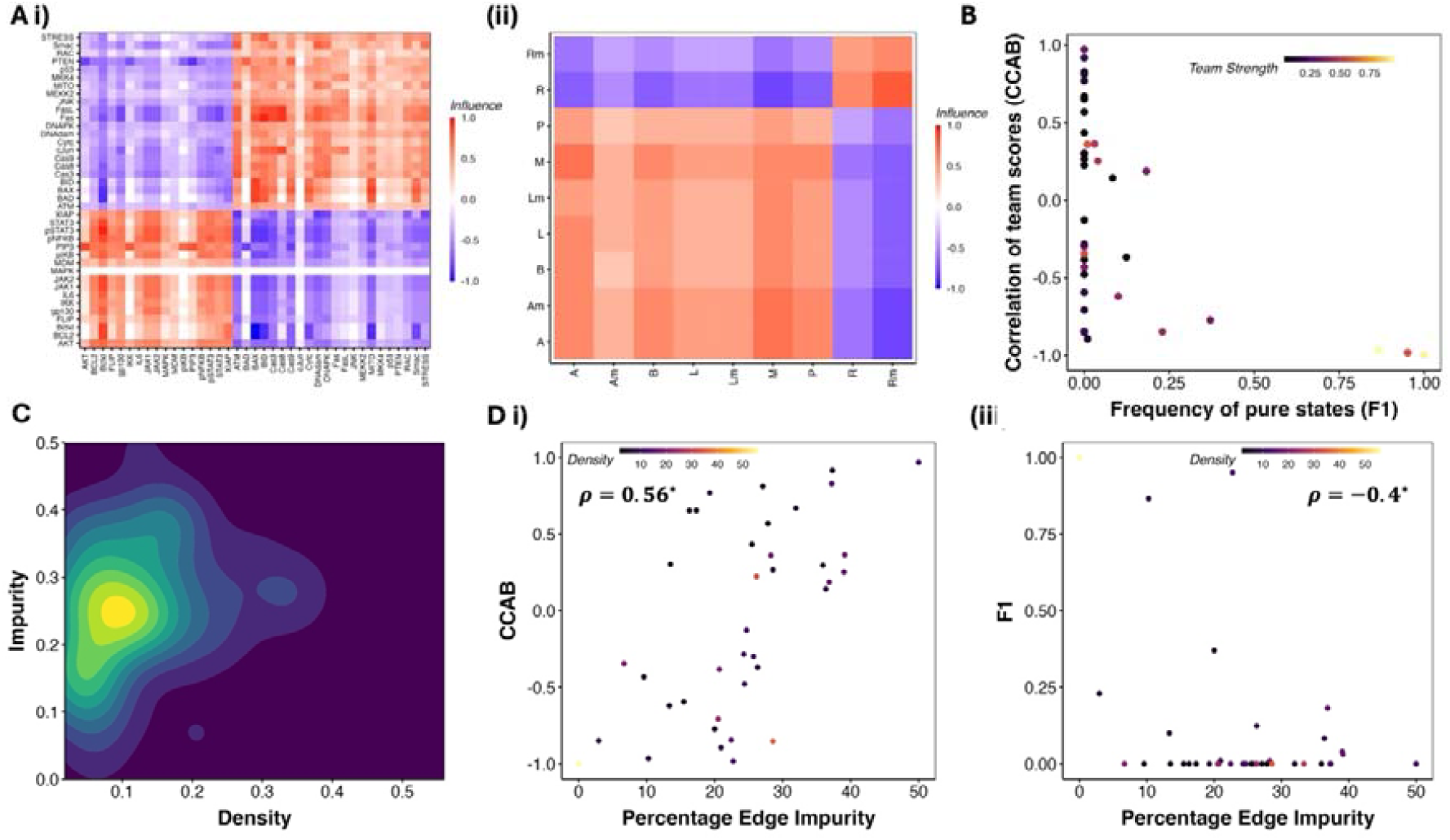
Team-based analysis of GRNS from the Cell Collective database. **A)** Examples of Cell collective GRNS with teams in mammalian (Chudasama et al. 2015) and bacterial (Veliz-Cuba and Stigler 2011). **B)** F1 and CCAB for cell collective networks obtained from simulating the logical rules. Each point corresponds to a network, colored by their team strength. **C)** 2-D density map of the biological networks depicting the Edge density and Edge impurity. **D)** Variations in CCAB and F1 with Edge Impurity.

To formalize this, we analyzed the *influence matrix* of each network, defined as the weighted sum of the powers of its adjacency matrix (see Methods). We then applied clustering algorithms to this matrix to classify the nodes into two provisional teams. Where needed, we refined these classifications manually to best approximate a mutually inhibitory team configuration.

For each network, we quantified two key structural metrics:

1. **Density**: defined as 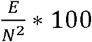 where E is the number of edges and N is the number of nodes in the network.
2. **Impurity**: defined as 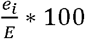, where e_*i*_ is the number of “impure” edges—those that are inhibitory between nodes within the same team or activatory between nodes in opposing teams. An impurity value of 0% corresponds to a perfect mutually inhibitory team structure, whereas an impurity of 100% indicates a configuration where nodes inhibit their team and activate the opposing team. Biological networks consist of some edges that may appear like impurities in the context of ‘team’ behaviour, such as auto-inhibitory feedback loops (for instance, seen in SNAIL1 (Peiró et al. 2006)) and incoherent feedforward loops (Lai et al. 2016), or mutual inhibition between EMT-TFs such as SNAI1 and SNAI2 (Subbalakshmi et al. 2022) or that between ZEB1 and ZEB2 (Vandamme et al. 2020). These structures play a crucial role in the contexts of robustness, noise dynamics, and signal transmission (Cinquin and Demongeot 2002; Dublanche et al. 2006; Zhang et al. 2012). Quantifying the effect of these inconsistent links can be challenging. The ‘teams’ conceptualization thus provides us with a way to quantify the extent of such inconsistencies through the lens of the impurity metric.

By incorporating the notion of impurity, we substantially relaxed the strict definition of a team, enabling a broader application of this concept across diverse GRNs.

To investigate the functional implications of these structural properties, we simulated the dynamics of each network using the logical rules provided in the Cell Collective database, under an asynchronous update scheme (see **Methods**). For each network, we initialized 100,000 random initial conditions and recorded the final steady states. For each such state, we calculated a “*team expression*” score, defined as the fraction of nodes within each team that were in the active (expression level 1) state. Similar to the cell fate decision networks earlier, the qualitative relationship between team strength, F1, and CCAB was maintained for cell collective GRNs as well (**Fig. 2B**). However, most of the networks had very low team strength, and in a low team strength regime, while F1 is consistently low, CCAB varied a lot, suggesting that team strength alone does not hold the differentiating power needed to predict the structure of the phenotypic landscape.

Across the Cell Collective GRNs, we observed an average network density of ∼15%, an average impurity of ∼30% (see **Fig. 2C**), and an average network size of 20 (**Fig. S3)**. Despite using the biological rules for simulation as opposed to threshold-based rules as in **Fig. X**, we found that the correlation between the expression levels of the two teams - CCAB was strongly positive with impurity and negative with density (**Fig. 2D i, Fig. S2**). Thus, impurity offers a better description of the phenotypic landscape than team strength. Networks with low impurity are closer in structure to two teams with mutual inhibition, leading to a strongly negative correlation between team scores. Whereas networks with a high impurity of (∼0.5) include the networks with mutually activating teams and therefore exhibit a strong positive correlation. Following our previous analysis of networks with teams, we calculated the frequency of “*pure states*” – F1, defined as the states with the team scores as 1,0, or 0,1. While the EMP networks showed a very high fraction of pure states (>75%, **Fig. 1C**), only three out of the 20 networks with teams in the Cell Collective database had high F1. F1 had a negative correlation with impurity (Spearman’s ρ = -0.4, p-value < 0.05) and a positive correlation with density (Spearman’s ρ = 0.31, p-value < 0.05). (**Fig. 2D ii**).

These findings suggest that a network’s structural characterization in terms of impurity and density can serve as a predictive indicator of its emergent phenotypic landscape. To probe this relationship further, we systematically generated artificial two-team networks (TTNs) spanning a range of impurity and density values and analyzed their dynamics using the same simulation framework. These results provide a unified framework to connect network architecture with phenotypic diversity, even in cases where traditional notions of well-separated teams break down.

### F1 and CC are strengthened with increasing density and team size, but weakened with increasing impurity

We generated artificial networks with two teams using the stochastic block model with constant probability for each community (Holland et al. 1983), as described in Fig. **3A**. Briefly, we generated an adjacency matrix of size 2N X 2N filled with zeros, where N is the number of nodes in a team. Each element of the adjacency matrix represents an edge from the node corresponding to the row to the node corresponding to the column. We then fill the matrix with 1’s (in the submatrices corresponding to within team interactions, i.e. pink) and -1’s (in the submatrices corresponding to across team interactions, i.e., yellow) in randomly chosen positions in the matrix, such that the total number of non-zero elements Is 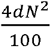, where d is the required percentage edge density of the network. We then flip the signs of a fraction 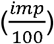 of the non-zero elements, where imp is the required percentage edge impurity. We generated an ensemble of the artificial two-team networks (TTNs), by varying 1) Team size (number of nodes in a team, changing as 5, 10 and 20), 2) Density (percentage of edges present in the network, ranges from 10% to 100%) and 3) Impurity (percentage of edges conflicting with the “team” configuration, ranges from 0% to 100%), with 100 networks for each combination of parameters, totalling 33000 networks. Note that we refer to density and impurity as “Core Edge Density” and “Core Edge Impurity”, where core edges refer to edges within the TTN. We use this terminology to differentiate the properties of the core network from the peripheral/external network later in the embedding simulations done later. described in **Fig. 5**. The range of team size was determined to capture the mean network size (20 – captured by team size 10) seen in cell collective networks

**Figure 3:**
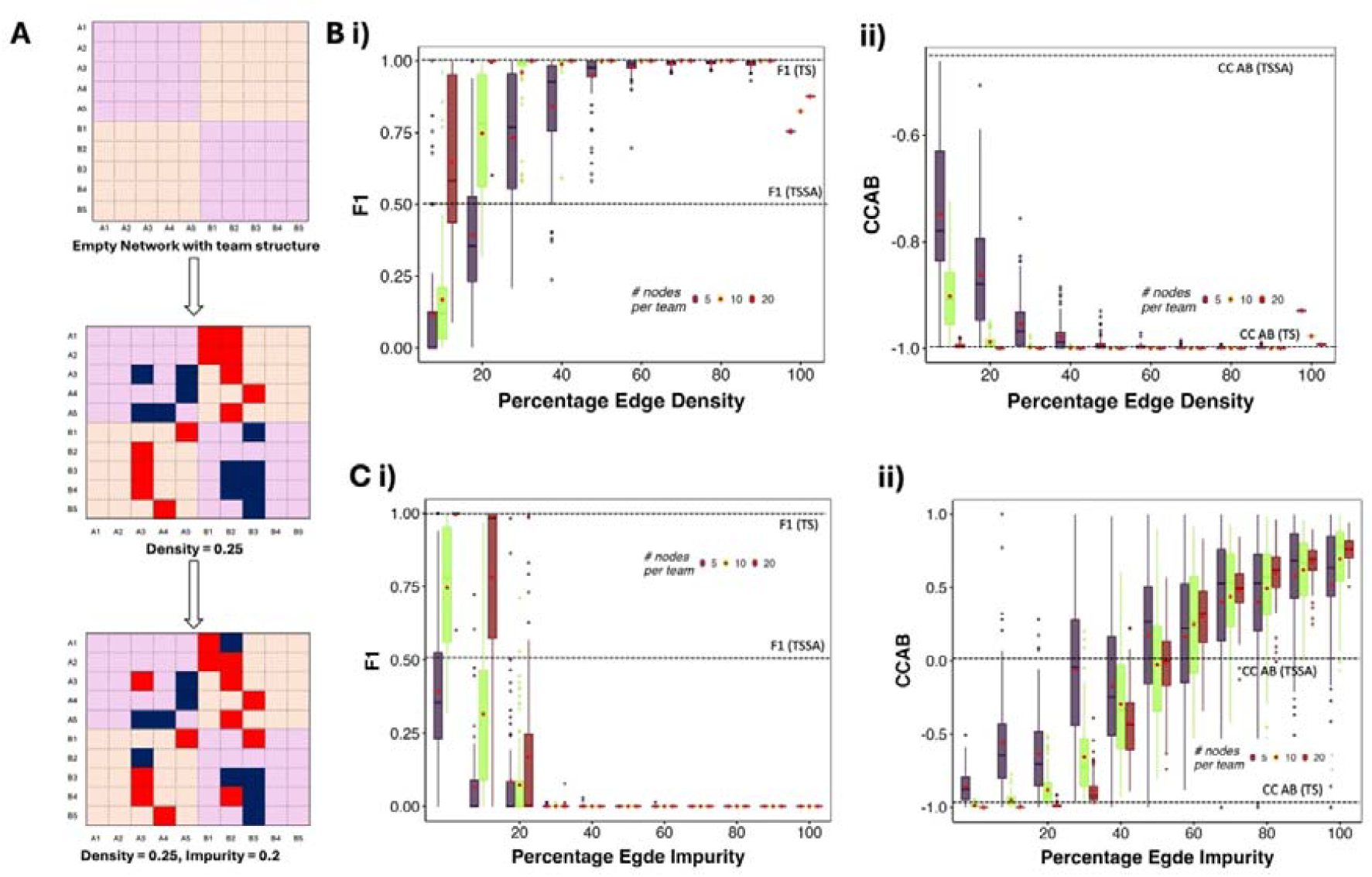
F1 and CC AB as a function of the edge density and impurity. **A)** Schematic representing the generation of TTNs. **B).** Boxplots showing the comparison between distributions of **i)** F1 and **ii)** CCAB with Variation in Percentage Edge Density at zero impurity. The boxes are colored by the team size, with the corresponding legend displayed inside the plot. **C)** Same as B, but variation against the Percentage Edge Impurity at a constant density of 20%. For each boxplot, the corresponding mean values of the metric on the y axis (F1 or CCAB) are shown using red diamond-shaped points.

We simulate the TTNs along with the toggle switch (TS) and toggle switch with self-activation (TSSA) networks using RACIPE (Huang et al. 2018) and Boolean formalisms (Font-Clos et al. 2018). RACIPE constructs and ODE system for a given network, then simulates it at a large ensemble of parameter sets to estimate the phenotypic landscape. The Boolean formalism used is a threshold-based formalism inspired by the ising model of ferromagnetic systems. Among various threshold formalisms available, the formalism used here mimics the Boolean networks in literature best (Kadelka and Hari 2025). From both formalisms, we obtain steady-state frequency distributions as a measure of the phenotypic landscape. We evaluated the bimodality of the phenotypic landscape using F1 - the total frequency of “pure” phenotypes (where all the nodes of one team have high expression and all those of the other team have low expression, equivalent to 10 or 01 states in TS) and the expression patterns using CCAB – the strength of correlation between the “teams” (see methods). We compared these against the calculations obtained from simulating TS and TSSA.

In the absence of any impurities (Hari et al. 2025), the frequency of ‘pure’ (mutually exclusive) states (F1) increases with increasing density and increasing team size in both Boolean and continuous simulations (**Fig. 3B, S4A, D**). While networks with 20 nodes per team saturated quickly, reaching F1 of 1 at 20% density, smaller networks required larger densities. The networks with 20 nodes showed a mean F1 of 0.75 at 20% density and saturated at 30% density. At 60% density, even the smaller networks reach an F1 of 1, the same as a toggle switch. The F1 of TSSA, believed to be the core network of many cell-fate decision systems, is 0.5. We find that most networks of team size 10 cross that threshold at 20% density, the mean density of cell collective networks. Team size 5 networks required 40% density to cross the threshold F1 of 0.5. This complementary relationship between team size and density could be the reason for the negative correlation between team size and density found in the cell collective.

In Boolean simulations, the maximum value of F1 is 1, and in continuous simulations, it is around 0.85. In both cases, the maximum F1 of TTN networks is equal to or above the F1 values obtained from a TS (as shown by the horizontal lines). Interestingly, at 100% density in both Boolean and continuous simulations, F1 values decreased from their maximum to approximately 0.8. This observation is analogous to the increase in the frequency of hybrid states in TSSA as compared to TS (Hari et al. 2022a, F1 in Fig. 3Bi). We hypothesize that up to 90% density, the activations within teams contribute towards making the teams work as single entities, beyond which the activation links within a team amount to adding a self-activation to the super-nodes, thereby increasing the occurrence of “hybrid” phenotypes. To further understand this behaviour, we removed all activation links from the TTN networks and found that F1 monotonically increased with density (**Fig. S4B, C)**. It is important to note that some networks at low densities and small team sizes also achieve very high values of F1 (see the outlier dots at 10% and 20% densities). Biological networks also showed similar behaviour at low density, where most of the networks have low F1, and a small fraction have higher F1.

The correlation coefficients of TTN networks remained strongly negative. At lower densities (<=20%), CC AB shows a higher variance. As the density increases, the variance decreases, with the mean approaching that of TS in both Boolean and continuous simulations **(Fig. 3B ii, S4C)**. Team size has a similar strengthening effect on correlation as density. Thus, TTN at any density can be simplified as a TS in the absence of impurities while studying the correlation between expression of teams/phenotypes. For the more nuanced studies involving the fraction of pure states in a population governed by these networks, tacitly simplifying TTN to TS or TSSA is not always valid. As biological networks have lower density, simplifying cell fate decision systems into simple toggle switches will lead to overestimation of pure states.

.Frustration of a steady state, a metric that measures the extent of inconsistency between the configuration of a steady state and the network topology, correlates negatively with the frequency of the steady state (Tripathi et al. 2020; Hari et al. 2022b). An activatory (inhibitory) edge is frustrated if the expression levels of the nodes corresponding to the edge do not have the same expression level (have the same expression level). Frustration of a state is defined as the fraction of such frustrated edges. In the absence of impurities, the ‘pure’ states (111110000 or 000011111 for a 10-node TTN) are not frustrated. However, as we introduce impurities in a TTN, the ‘pure states’ of the original TTNs become frustrated, with the value of frustration equal to the fraction of impure edges in the network. Given the previously observed negative correlation between frustration and steady state frequency (Hari et al. 2022b), we predict the F1 metric to decrease as the number of impurities increases in the network. **Indeed, we find that impurity has a negative effect on F1 at different team sizes (Fig. 3C i)**. The mean F1 for all network sizes quickly reduced to zero as impurity increased to 30%. Even at 80% density, where networks of all sizes had an F1 of 1 at 0% impurity, the F1 decreased to 0 at 40% impurity **(Fig. S5A)**, demonstrating the strong impact impurity has on the TTN phenotypic landscape. It is interesting to note that the average impurity of biological networks is 25%, and the density is around 20%. Thus, our results align with the low F1 values found in cell collective networks, despite the difference in simulation formalism. To better understand the decrease in F1, we studied additional metrics that relaxed the definition of F1 by including the frequency of states with deviation from pure states up to one and two nodes (**Fig. S5B)**, thereby creating two relaxed F1 metrics. While the mean values of these relaxed F1 metrics are larger than those of the strict F1, we still see a sudden decrease in the mean values of these metrics from 20% to 40% impurity. A subset of networks do show higher values of the relaxed metrics as well as the strict F1, suggesting the existence of additional network features that can regulate the effect of impurity on F1. From 60% impurity onwards, both the relaxed F1 metrics have zero values as well.

CC AB provides a better understanding of the changes in the phenotypic landscape as compared to F1 with changing impurity. The mean correlation strength between the two teams reduces as impurities are introduced, nearing zero as the impurities reach 50%. With further increase in impurities, the correlation coefficient changes sign from negative to positive and starts increasing, reaching 1 at 100% impurity **(Fig. 3C ii, S5C)**. At 100% impurity, the TTN is effectively no longer a mutually inhibiting teams’ case, but a mutually activating one, leading to strongly positive CCAB. Correspondingly, we see the frequency of states with both teams “on” or both teams “off” increase with higher impurity **(Fig. S5D)**. Similar to F1, at 20% impurity, more than 75% of the networks show a stronger correlation than TSSA, but most networks have a weaker correlation than TS (-1). At 60% impurity, a significant fraction of the networks (>25%) still showed negative CC AB. While at 80% and 100% impurity, when the edges with each team are mostly inhibiting and those across teams are mostly activating, we still see a small number of networks showing negative CC AB.

### Impurities have a stronger effect on the phenotypic landscape than density and team size

After analysing the effect of density and impurity individually, we investigated the combined effect of different impurity and density values on F1 and CC of the TTN networks. At all densities, the networks with 60% or higher impurities show an F1 value of 0, indicating that high impurity can impair the TTN functioning irrespective of density. At 20% impurity, we observe a non-zero mean F1 for all densities as seen earlier (compare **Fig. 4A with Fig. 3C)**. As density increases, we see an increase in the mean value of F1 for networks with 20% impurity, indicating that density can make the phenotypic output resilient to these impurities. Note that the magnitude of the increase in mean F1 is lower, from 80% to 100%, as compared to other consecutive density points. However, at 100% density, the mean F1 for 20% impurity networks is very similar as that at 0% impurity, while there is a strong decrease in mean F1 for other densities (**Fig. 4B)**. This observation indicates that high density prioritizes negating the effects of impurity over adding self-activation to the network. At all other values of density, we see a clear decrease in mean F1 as impurity increases (**Fig. 4B**). The median values of F1 (horizontal dashes within the boxplots) follow the same trend as that of the mean. First, we take a closer look at the networks with 20% impurity in **Fig. 4B** (orange boxes). The range of F1 values is 0 to 1 for densities 40% to 100%. At 20 and 40% density, a large fraction of the networks have low F1 values. The span of the inter-quartile range (IQR) for F1 decreases from 40% to 100% density. However, the upper limit of the IQR increases from 40 to 80% density, but goes back down at 100% density, reminiscent of the decrease observed in **Fig. 3B**. The lower limit of the IQR consistently increases with an increase in the density. We interpret this behaviour in the following manner. On average, an increase in density increases F1 while an increase in impurity reduces F1. Near the lower limit of the IQR, impurity dominates, while near the upper limit, density dominates, evident by the decrease in F1 at 100% density as compared to 80% density. However, other network features can impact this visible competition between impurity and density.

**Figure 4:**
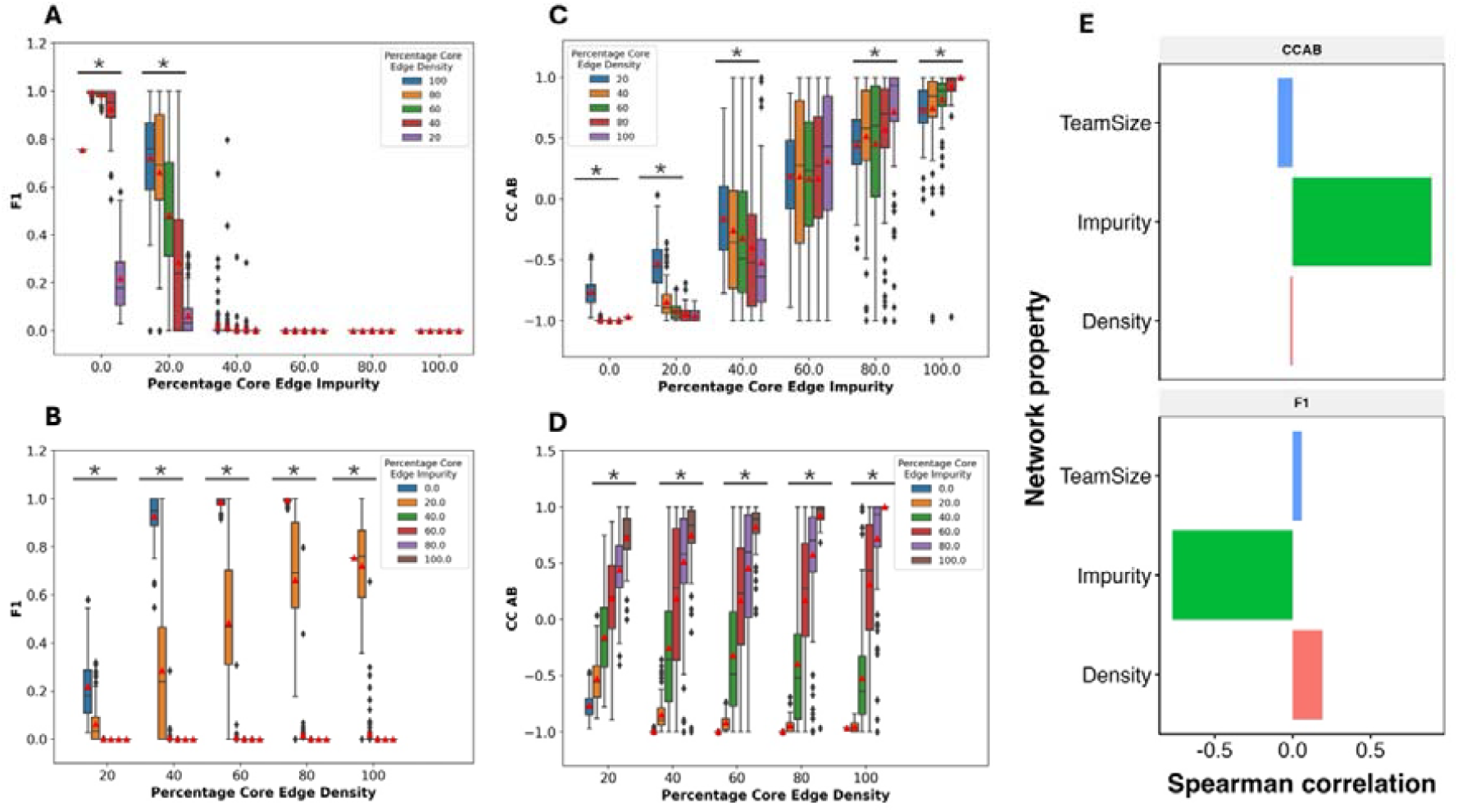
Varying the percentage of impure edges and density together. Comparison between the distributions of A) F1 and C) CC AB for TTNs of team size 5, having the same Percentage Core Edge Impurities but different Percentage Edge Densities. Comparison between the distributions of B) F1 and D) CC AB for TTS-5 networks, having the same Percentage Core Edge Density but different Percentage Edge Impurities. ^*^ indicates that the p-value of one-way ANOVA is < 0.05. (E) Bar plot depicting the Pearson’s correlation coefficient of Network parameters, percentage core edge density, and impurity with F1, CCAB, and Team size.

The correlation coefficient reveals further interesting observations in this regard. As seen in **Fig. 3C**, the correlation coefficient moves from value close to -1 to a value close to 1 as impurity increases, following a shift in the network structure from mutually inhibiting teams to mutually activating teams **(Fig. 4C)**. As the density increases, the range of CC values over the same range of impurity values increases **(Fig. 4D)**. We found similar dependence of F1 and CC AB on density and impurity for team size of 10 **(Fig. S6)**. When density is varied at constant values of impurity, the direction of change in correlation coefficient depends on the impurity value **(Fig. 3B)**. For impurity less than 50%, we see a decrease in the correlation coefficient (i.e., CC becomes more negative), whereas after 50%, CC increases, becoming more positive. Note that below 50% impurity, mutual inhibition of teams is dominant, while above 50%, mutual activation is dominant, possibly explaining the above-mentioned trend. Overall, we find that the increase in density amplifies the impact of the dominant behaviour of the network in terms of the correlation coefficient. Thus, unlike in cases of no impurity (**Fig. 3C**), we see a clear effect of density on CC in the presence of impurities.

While F1 is a strict measure (**Fig. S5)**, CC AB is more flexible. At 40% impurity, we do not see any networks with an F1 of 1, and a very low fraction of networks with non-zero F1. However, at 40% and 60% impurity values, we see networks with 1 and -1 correlation coefficients at all densities above 20%. While the changes in mean F1 and CC are captured well by the density and impurity of the TTN networks, the wide range of values suggests that predicting these metrics for individual networks would require additional understanding of the network topology.

To calculate the relative effect of density and impurity on F1 and CC, we calculated pairwise correlation between the density and the impurity vectors with F1 and CCAB **(Fig. 4E)** vectors. In other words, we measured the association of each parameter with F1 and CCAB as the other parameter(s) is varied over a wide range. As expected, impurity has a negative (positive) correlation with F1 (CC AB) while density has a positive (negative) correlation with these metrics correspondingly. We find that impurity has a stronger correlation with both F1 and CC as compared to density, suggesting that impurity has a stronger impact on both these metrics. Because F1 falls to zero after 40% impurity, we studied the networks with impurity 40% separately and found that impurity still has a stronger correlation with both F1 and CCAB than density does. At a constant impurity, the effect of density on CCAB is much lower **(Fig. 3C)** as compared to the effect of impurity at a constant density **(Fig. 3D)**.

### Resilience to global perturbations – interaction of teame-based networks with large random networks

One factor often ignored in GRN-based models is the interaction of these networks with the rest of the signalling networks within which they are embedded. Indeed, biological networks have been argued to be modular (Serban 2020; Alcalá-Corona et al. 2021; Kadelka et al. 2023), but multiple phenotypic plasticity programs are interlinked, such as EMT, stemness, and drug resistance through a few intermediary molecules and pathways (Pasani et al. 2020; Bangarh et al. 2024). Thus, any network reduction strategy in terms of identifying teams of nodes should be capable of recapitulating the effect of such global interactions. We had previously shown that: 1) Toggle switch motifs are resilient to perturbations from such “global” interactions to a large extent (Harlapur et al. 2022). 2) The correlation and PC1 variance of larger networks are significantly affected by the presence of a sub-network with teams (Hari et al. 2025). These observations indicate that teams can provide resilience to global interactions similar to that seen in a toggle switch. Thus, we investigated the effect of global interactions on the phenotypic outcome of TTN.

We embedded TTNs in a larger external network of 30 nodes, sampling external network parameters from the ranges mentioned in **Table 1** (see schematic in **Fig. S8**). We investigated three external network parameters that can potentially influence the dynamics of the core network: 1) Mean Connectivity, a measure similar to density that describes the number of connections each node can have within a network on average. 2) Average indegree to the core node, measured as the average number of connections from the external network to the nodes of the core network, and 3) Average outdegree of the core node, measured as the number of edges from the nodes of the core network that can affect the core network dynamics via feedback through the external network and the indegree to the core network. We first generated the core and external networks separately. The core network was generated using the stochastic block model as described earlier, varying the core edge density, core edge impurity, and team size as required for the particular experiments **(Fig. S8A)**. For the external network, we generated a matrix of dimensions 30 X 30 filled with zeros, where each cell represents an edge from the corresponding node in the row to that in the column. We then randomly chose *Mean Connectivity* * 30 edges in the matrix and assigned 1 (activation) or -1 (inhibition) to these edges with equal probability (**Fig. S8B)**. We then combined the core and external matrices as shown in **Fig. S8C**. We then added *N* * *Average Indegee* edges (randomly choosing activations and inhibitions with equal probability) to the top right rectangle that contains the edges from the external network to the core network and *N* * *Average Indegee* edges to the bottom left rectangle that contains the edges from the core network to the external network, where N is the number of nodes in the network **(Fig. S8D, E)**. Note that the mean connectivity relates to the density of the external network as follows:

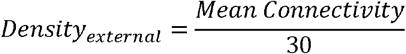

**Table 1.**
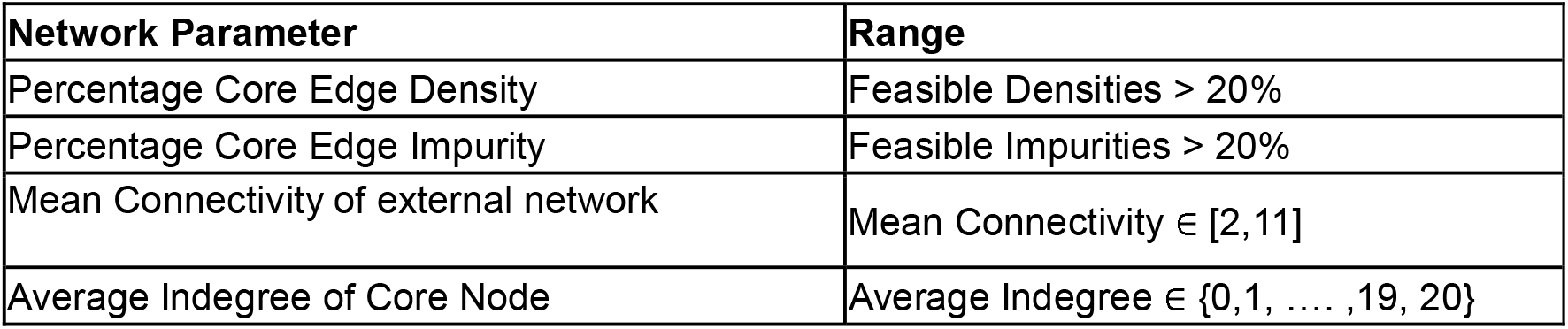
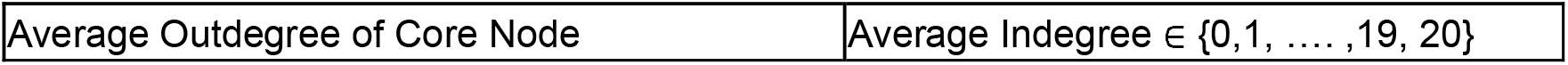
Ranges for different parameters considered for network embedding

We refer to the mean connectivity instead of density for the external network so that it remains distinct from core edge density. The choice for the range of mean connectivity was inspired by the distributions of connectivity reported for an ensemble of biological networks (Kadelka et al. 2024). We first wanted to test the effect of core network parameters on F1 of an embedded TTN, starting with the team size. We generated 100 such networks for each value of core team size (varying from 1 to 6) with 100% core edge density, 0% core edge impurity, and chose the external network parameters randomly from the ranges described in **Table 1**. Apart from team size 2, TTN shows higher mean F1 values than the toggle switch when embedded in a larger network **(Fig. 5A**, compare team size 1 with higher team sizes). The mean F1 saturates at 1 for team sizes above 5, suggesting that larger TTN networks are more resilient to external perturbations. This inference was reinforced in CC AB, where the correlation remained strongly negative from team size 2 onwards **(Fig. S9A)**. Based on this saturating behaviour and computational costs of simulations, we chose a Team Size of 5 for further embedding experiments, such that we are simulating 40 (= 10 + 30) node networks.

**Figure 5:**
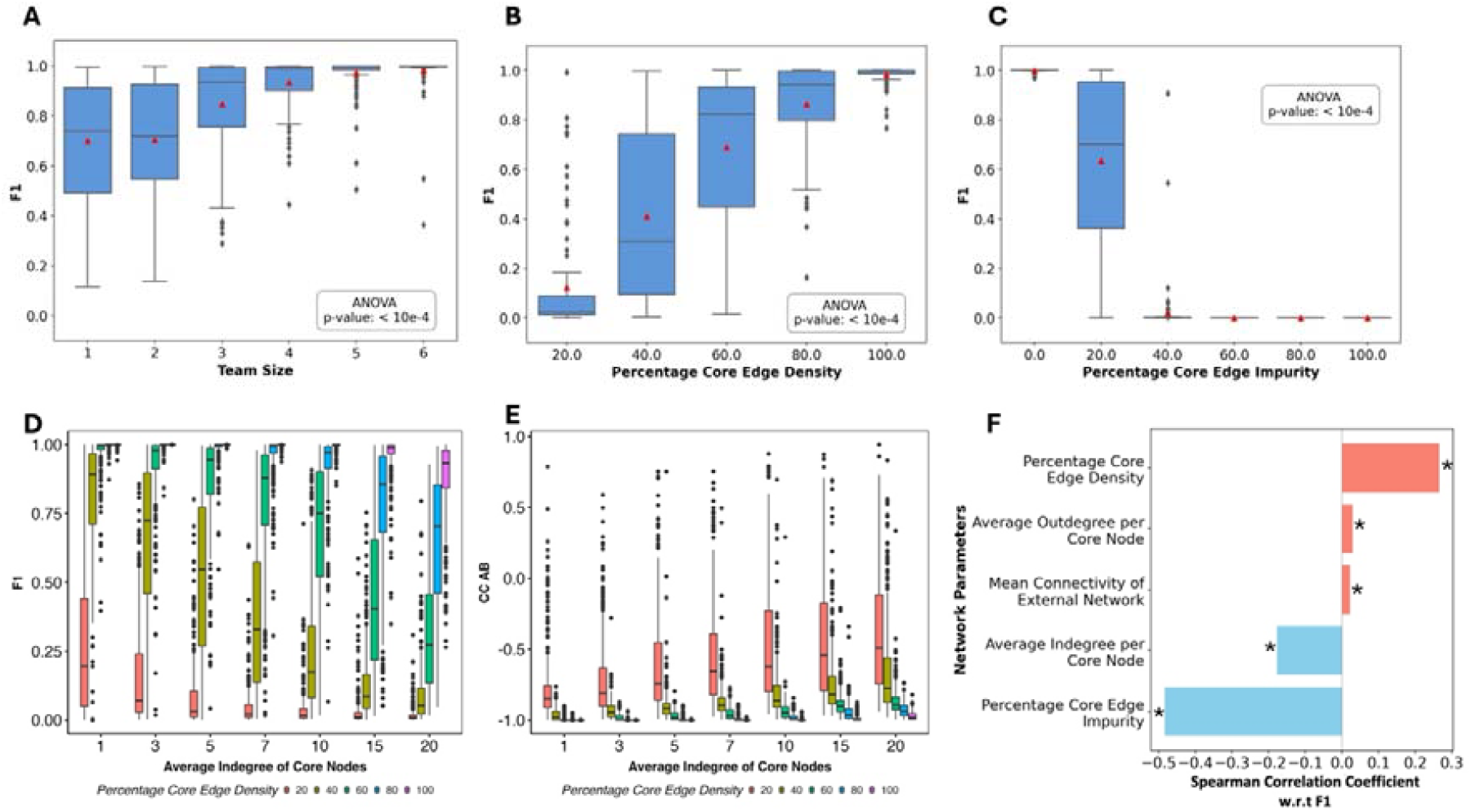
Dynamics of Embedded Teamed Toggle Switch Networks. **Boxplots comparing the Distributions of F1** for variation in A) Team Size, B) Percentage Core Edge Density, C) Percentage Core Edge Impurity, and D) Average Indegree per Node of a Team of fully connected TTN-5 networks embedded in a 30 Gene External networks. Bar plot showing the Pearson’s correlation coefficient of F1 with Percentage Core Edge Density, Impurity, Mean Connectivity of External Network, Average Indegree per Core Node, and Average Outdegree per Core Node. ^*^ Indicates p-value < 0.05.

Next, we varied the density of the embedded TTN, keeping the size at 5 nodes per team and impurity at 0%, and randomly choosing the external network parameters from their corresponding ranges. Similar to the unembedded case, the mean F1 increases with increasing density. While the mean F1 reaches 1 around 60% density in the unembedded case, it takes 80% density for the mean F1 to reach 1 in embedded networks **(Figure 5B)**. Team correlation remained negative for all densities **(Fig. S9B)**. Interestingly, TTN could only withstand up to an impurity of 20% before losing the F1 frequency, suggesting that internal perturbations affect the phenotypic landscape more than external perturbations. We similarly explored the effects of core impurity on the embedded TTN landscape (**Fig. 5C)**. As expected, increased impurity reduces F1, with mean F1 falling to 0 at 40% impurity. At 20% impurity, the variation in F1 increases upon embedding (compare **Fig. 2D** with **Fig. 5C)**, with more networks having a lower value of F1 due to the combined effect of internal and external perturbations to TTN. Similar to the unembedded case, the CC AB values increased with increasing impurity, turning positive at impurities higher than 60% (**Fig. S6C)**.

We then explored the resilience that TTN networks have to external perturbations by controlling for the variables describing how well the external network is connected to the core network. Previously, we had found the toggle switch to be resilient up to an in-degree of 5 on each of the two nodes. Here, we defined the indegree of each node as the average indegree of the nodes of the TTN (for example, if the total number of incoming edges on TTN is 10 for 10 nodes in TTN, the indegree is 1). Note that this definition implies that the indegree of a team from the external network is N times the average indegree of each node, where N is the number of nodes in a team (5 here). We generated TTN embeddings for multiple combinations of values of in-degree and density. For each of these networks, we maintained the mean connectivity of the external networks at 11 to ensure the maximum impact while randomly sampling the average outdegree of core nodes. Interestingly, we find the F1 for TTN at 100% density is resilient to large values of in degrees, with the values of F1 staying at 1 up to in degrees of 10, and mean F1 reaching 0.85 at in degrees 20 **(Fig. 5D, pink box)**. At lower densities, the effect of indegree could be seen on the decreasing F1 value, as expected. However, TTN proves to be more resilient than a toggle switch to external perturbations even at lower densities. While the toggle switch lost the features of bimodality after an indegree of 5, TTN maintained their F1 at a team indegree of 5 (compare average indegree of 1 in **Fig. 5D** with **Fig. 3E**). A similar effect of indegree and density could be seen (**Fig. 5E)** where, as indegree increases, the CC AB increases. Note, however, that CC AB remains negative in all cases, except for a small fraction of networks at 20% density. These results show that the TTN is highly resilient to external perturbations in the absence of impurities.

To better understand the dependence of F1 on the various network parameters studied in this section, we generated 2500 networks with random values of the parameters sampled from the ranges described in **Table 1**, and calculated the F1 values emergent from each network. We then calculated the correlation coefficient between each parameter and the F1 vectors as a measure of the relative dependence of F1 on the metrics. We found that F1 depends strongly on the core impurity, followed by core density, evident from the strength of correlation between the vectors **(Fig. 5F)**. Overall, our results suggest that TTN is highly resilient to external perturbations, making its study as an isolated model valid in the presence of the global network. Furthermore, TTN has a higher resilience than a toggle switch to external perturbations.

#### Characterizing the phenotypic landscape as a function of Density, Impurity, and team size

Our analysis so far has evaluated the phenotypic landscape from the point of view of its potential to be a bimodal landscape with two mutually exclusive states as the two dominant phenotypes. We further evaluated the correlation between the team expressions as a primer towards understanding the expression trends of the defined teams in the network. While these metrics serve well to understand how well a given network gives rise to a bimodal phenotypic landscape with mutually exclusive expression patterns in the dominant phenotypes, as observed in binary cell fate decision networks. Thus, our analysis so far sets the limits under which a network can act as a binary cell-fate decision network: namely, an impurity less than 30% and either a large network size or a reasonably high density, ensuring ample connectivity. However, these metrics are clearly lacking in providing a complete description of the phenotypic landscape, and are therefore not enough to evaluate the utility of a team-based view of GRNs to predict the properties of the phenotypic landscape. For example, at high impurity values (> 40%), we know that F1 goes to zero, and the correlation between the team scores starts to become positive. But what kind of states show up in the absence of pure states? In this section, we will focus on answering such questions using appropriate metrics. Through this analysis, we aim to evaluate whether the team’s perspective is useful to understand the behavior of any GRN.

In line with our goal to maintain a team-based approach to the phenotypic landscape, we used a team score-based representation for each steady state. Each steady state is represented by two numbers, each number less than one representing the fraction of nodes in one of the teams that have an expression level of 1. Thus, the pure states become either (1,0) or (0,1). We then calculated the mean frequency of such team configurations for each impurity, density, and team size, and asked if any obvious patterns in steady states emerge (**Fig. 6A, S10)**. At low impurity (=< 20%), the pure states dominate. At very high impurities (>= 80%), the highest frequency was occupied by states around (0.5, 0.5), in the direction of T1 = T2, as indicated earlier by the positive CCAB at large values of impurities. At all the intermediate levels, the mean frequency remained largely uniform across all the team expression compositions. To further understand this behavior better, we calculate the “balance” of each steady state, defined as a product of scaled team scores, as follows:

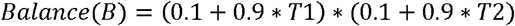

**Figure 6:**
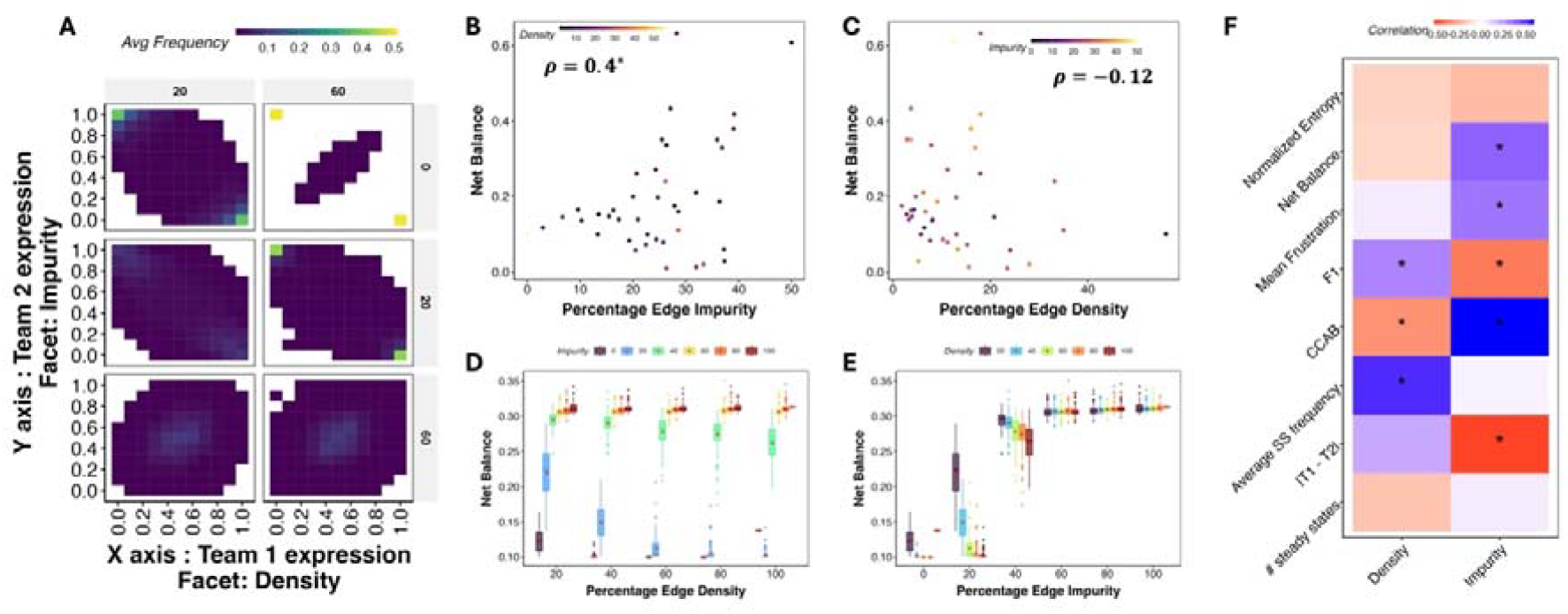
Phenotypic landscape of GRNs as a function of density and impurity. A) Average frequency of different combinations of team scores at steady state for different density and impurity values. B-C) Net balance as a function of Impurity and Density for GRNs of cell collective networks. D-E) same as B-C but fo artificial networks. F) Correlation between Density, Impurity and metrics of the phenotypic landscape for Cell Collective GRNs

Where T1 and T2 are the scores of team 1 and team 2, respectively, for each given state. We then define the net balance for a network as the weighted sum of the balance of all steady states, as follows:

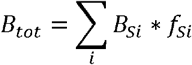

Where *Si* is a steady state, *f*_*Si*_ is the frequency of the steady state. B ranges between 0 and 1. Note that each value of B uniquely identifies a pair of T1 and T2. Furthermore, a high B is only possible when both T1 and T2 are high, whereas a low value can be obtained by at least one of the team expressions being low. Thus, as a rule of thumb, states with highly unbalanced configurations, such as the pure states, end up having low values of balance, while states with highly balanced configurations (i.e., similar values of T1 and T2) have relatively higher values of balance.

In cell collective networks, we found a positive correlation between balance and impurity and a negative correlation between balance and density (**Fig. 6B, C)**. We found similar relationships in the artificial networks as well. Balance increases as impurity increases, and decreases as density increases **(Fig. 6D, E)**. Interestingly, the mean of balance saturates at around 0.3 after an impurity of 50%, which corresponds to a team score of nearly 0.5 in both cases. From the heatmaps in (**Fig. 6A and S10)**, we find that this saturation coincides with a uniformly low expression of all team score configurations. To confirm, we studied how the difference in team scores changes with Impurity and Density, and found it to be consistently low at. Thus, together these results suggest that after an impurity of 50%, it may be unrealistic to try an predict the state configuration using the teams architecture. There is also a significant mismatch in the balance values between the cell collective and artificial networks. Generally speaking, cell collective networks seem to have a higher balance value than the artificial networks. One possible reason for this behavior could be that cell collective networks contain many special cases, such as networks with single teams (der Heyde et al. 2014), where the influence matrix does not contain any negative values. The steady states of this network will indeed have high balance, due to the strong mutual activation pushing all nodes to be active or inactive together. In our framework, this network will be classified as having 50% impurity. However, random network generation cannot hope to achieve such an extreme configuration, and even if it does, it will end up as an outlier of the distribution.

We calculated additional metrics to further gain insights into the phenotypic landscape **(Fig. S11)**. As impurity increased, the number of steady states initially showed an increasing trend till 40%, then decreased till 100%. At low impurity, high density led to reduced entropy, but at higher impurity, density had a non-monotonic effect as well. This can be explained by thinking of impurity as the primary effector, with density supporting whichever direction the impurity takes the network in. At low impurity, mutually inhibiting teams dominate, while at high impurity, mutual activation dominates. Correspondingly, good control is exerted on these extreme cases. Similar results were observed for entropy as well. The frustration of the steady states monotonically increases with impurity, while the average frequency decreases. In line with the analysis of artificial networks, we found strong, significant correlations for Cell Collective GRNs between Impurity and metrics of phenotypic landscape: frustration, net balance and the difference in team scores across all steady states of all networks, in addition to F1 and CCAB (**Fig. 6F**). Overall, our results provide trends to predict the phenotypic landscape based on the density and impurity, and find that moderate impurity leads to high entropy, high frustration, and low average frequency, and in general reduces the determinism of the dynamics. But at low and high impurity, we can predict the dynamics of any GRN with reasonable accuracy.

## Discussion

Teams are a topological feature of binary cell-fate decision networks that we have observed in multiple biological contexts (Chauhan et al. 2021; Pillai and Jolly 2021; Shyam et al. 2023; Sehgal et al. 2024; Hari et al. 2025). The “influence matrix” representation of these networks had revealed two communities that are identified by groups of nodes with positive influence within themselves and negative influence across, forming a “Two-teamed network” (TTN) (Chauhan et al. 2021; Hari et al. 2022b). This influence is mediated through both direct and indirect interactions between each pair of nodes. An intuitive coarse-graining of the influence matrix leads to a toggle switch of two supernodes, with each supernode representing one team **(Fig. 1B)**. However, we find here that such coarse-graining of these networks is not always appropriate.

We studied three key features of the TTN: the size of each team, the density of the network, and impure edges that have the opposite nature as appropriate for the team classification (i.e., inhibitions within teams and activations across teams). The characteristic features of a toggle switch are: two dominant steady states (10, 01) with bistability and consequently a strong negative correlation. Consequently, we used the total frequency of pure states (expression of nodes belonging to one team is on, while that of the other team is off) and the average correlation across nodes of the network as representatives of the phenotypic landscape of TTNs. While the correlation coefficient is a low-resolution measure of the mutual inhibition between the teams, F1 is a more nuanced and strict metric describing how often these networks can give rise to, as opposed to two dominant cell fates that the network is supposed to give rise to.

Our analysis revealed that in terms of F1 and CCAB, TTN networks needed to have high density/network size and low impurity to perform as well as a toggle switch. However, TTNs overtook the TSSA at reasonably low values of density and maintained better F1 and CCAB till higher values of impurity. But our analysis also revealed several interesting relationships between the network structure and the metrics of the phenotypic landscape. At low impurity, TTNs gave rise to bimodal phenotypic landscapes indicated by high F1 and strongly negative CCAB. As impurity increased, F1 quickly dropped while CCAB rose from -1 to 0 at 50% and 1 at 100% impurity. Previously, our quantification of team strength failed to predict the structure of the phenotypic landscape at low values of team strength. Including impurity as a metric is an improvement in this direction, evidenced by the relationship between CCAB and impurity.

The usefulness of impurity is further supported by our analysis of cell collective networks. We analyzed 42 GRNs from the Cell Collective database (Helikar et al. 2012), simulating them using their corresponding logical rules. We coerced these networks into a two-team format, calculating their impurities. We found that network impurities correlated strongly positively with CCAB for cell collective networks, suggesting that the idea of using teams to understand network behavior is indeed useful. While studies that seek to understand the role of network structure either focus on smaller-scale properties such as feedback loops applicable to smaller networks or statistical metrics such as connectivity and spectral characteristics applicable to larger networks, teams provide a balance between the two ways by both applying to larger networks while providing a meaningful way to incorporate feedback loops. Low impurity is an indicator of cohesive positive feedback within the network, while higher impurity is an indicator of disruptive negative feedback. This is evident from the frequency of team score combinations **(Fig. S10)**. While at low impurity, node teams cohesively oppose each other, at moderate to high impurity values, the landscape is diluted, with a significant increase in the number of steady states, entropy, and decreasing predictability. At very high impurity, on the other hand, we found the state space concentrated around partial expression of both teams together, a behavior reminiscent of hybrid states.

Other interesting observations enhanced our insight into the network structure-phenotype connection. At 100% density and 0% impurity, F1 goes down to 0.8, while at 90% density it is at 1. This behavior is similar to an increase in the frequency of the 11 and 00 states in TSSA as compared to TS. Indeed, removing all the within-team interactions, thereby reducing the self-activation on the supernodes, increased F1 back to 1. On the other extreme, at 100% impurity, the strong positive correlation was interesting, since 100% impurity implies positive interactions across teams and negative interactions within. Together, these results highlight that interactions across teams have a higher impact on the phenotypic landscape than those within teams.

The internal properties, such as density and impurity, were able to explain the trends of mean F1 and CC well, but failed to explain the occurrence of some exceptions, especially in the presence of impurity. Even at 100% impurity, we found some networks with a team correlation of -1. Similarly, at 40% impurity, where the interactions are nearly random, we find networks with high values of F1. Explaining such apparent anomalies would potentially invoke including additional topological features in the network such as node connectivity measures (Wang et al. 2022), strongly connected components, stable motifs (Zañudo and Albert 2013) and network redundancy (Gates et al. 2021).

Our analysis of network embedding further provides key insights into how cell-fate decision systems can be dynamically modular (Deritei et al. 2016). Even at lower densities, we found that the dynamics of TTN are resilient to the influence of the external network. Together with our previous results on the robust low-dimensionality of TTNs, we propose teams as an underlying mechanism for modular dynamics of phenotypic switches, such that they maintain their phenotypes when interacting with other switches/signalling networks in the cell.

We further studied the configuration of all the steady states, trying to get a holistic picture of the effect of density and impurity on the phenotypic landscape. Using a metric we termed the balance of the phenotypic landscape, we found that the loss of F1 with an increase in impurity does not lead to a gain of any other specific states. Even at high impurity, where the correlation turned completely positive, there was no consistency in the steady-state configurations. This behavior can be explained by the fact that mutually inhibiting teams exert a strong control over the phenotypic landscape. The impurities that disrupt this control, while enabling some general coexpression-like properties, do not control the phenotypic landscape well, leading to what can be thought of as a continuum of states.

We acknowledge several important limitations. Direct experimental validation of the relationship between impurity and phenotypic outcomes remains limited due to the scarcity of comprehensive datasets linking network topology to phenotypic landscapes. Inference of large-scale networks that can capture team effects is constrained by insufficient time-course data, and currently available inference methods lack reliability and applicability across diverse datasets (Pratapa et al. 2020). Our Cell Collective analysis represents a pragmatic approach to analyze networks that have been individually constructed from and validated against biological data, providing the best available foundation for testing our theoretical predictions. Future experimental work using single-cell trajectory analysis, systematic perturbation studies, and synthetic biology approaches with controlled network architectures will be essential to validate these findings in living systems. We made multiple assumptions in our simulations, which would require careful consideration in future studies. First, we simulated the artificial networks mainly using a threshold-based Boolean formalism. While threshold-based formalisms that are often employed in modelling Gene regulatory networks (Mendoza and Alvarez-Buylla 1998; Li et al. 2004; Font-Clos et al. 2018) cannot account for the logical rules and interaction parameters that govern the emergent dynamics (Zañudo et al. 2011). Second, we assumed that both teams have equal sizes. While this is not an issue with the threshold-based formalism used to simulate the networks here, a different simulation formalism might lead to asymmetry in the phenotypic landscape caused by the difference in team sizes. However, this might not matter for F1, as the metric measures the extent of pure states and does not distinguish between the pure states that emergent from the system. Similar symmetry was maintained while constructing the networks, making sure that each of the four submatrices of the interaction matrix defined by the team structure has near-identical properties. Investigating the effect of these asymmetries on the phenotypic landscape will be the focus of future studies. We also only considered features related to teams to try and explain the phenotypic landscape. While by design, this study to aims to understand how successful this exercise can be, the fact that impurity is unable to explain various aspects, such as the outlier networks, the configuration of phenotypes at intermediate impurity and lower density regions suggests that a complete picture of the structure function relationship in GRNs would require including additional structural metrics.

Network coarse-graining is a common strategy when dealing with large-scale networks, often involving the replacement of a group of nodes with a single “super” node, based on features such as positive feedback and strong connectedness, and communities (Erban et al. 2006; Saadatpour et al. 2013; Snowden et al. 2017; Caranica and Lu 2023; Kadelka et al. 2023). Inferring gene regulatory networks from perturbation time-course experimental data also allows identifying such coarse-grained structures (Wang et al. 2009; Sarmah et al. 2022). Binary cell fate decision systems are often modelled using toggle switches, serving as coarse-grained approximations of much larger networks comprising numerous transcription factors corresponding to each of the two cell fates (Roeder and Glauche 2006; Zhou and Huang 2011; Okawa et al. 2016). Indeed, simulations of corresponding large-scale networks, wherever available, do support the approximation as a toggle switch qualitatively in terms of predominant phenotypes (Ríos et al. 2015; Udyavar et al. 2017; Font-Clos et al. 2018). However, studies pertaining to the topological features of large networks that allow for a quantitatively valid coarse-graining of large-scale networks are missing in the current literature. Furthermore, a quantitative coarse-graining of larger networks into smaller networks is not always feasible, as demonstrated in our analysis of TTNs. Our view of teams with impurities applied to different GRNs can help with their dynamical coarse-graining. For example, one could develop and ODE model of a toggle switch to represent the larger networks, with noise in the parameters corresponding to the interaction between nodes that depends on the impurity within the larger network. This exercise remains to be of interest for our future investigations.

Overall, our results provide a theoretical investigation of topological features observed widely in cell-fate decision-making GRNs, and their applicability to representing and understanding GRNs that do not directly fit the confines of binary cell-fate decisions. The effect of such topological range on metrics such as correlation and entropy, which are commonly studied in high-throughput data, demonstrates the effect network topology can have on the emergent behavior such as co-expression, and can help provide an estimate of the range of network topologies that can underly high-throughput data and in turn can inform the predictions into dynamics meaningfully.

## Methods

### Identification of teams in GRNs

We first calculate the influence matrix (Hari et al. 2022b) using the following formula:

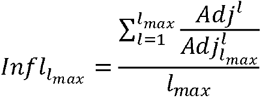

Where *Adj* is the adjacency matrix of the GRN, with each element of the matrix representing the sign of the edge, either -1, 1, or 0, from the node on the row to the node on the column. *Adj* _*max*_ is the unsigned form of the adjacency matrix. *l*_*max*_ is the maximum path length for which influence is calculated, typically 10. We then apply hierarchical clustering on the 2*N X N* matrix (where N is the number of nodes) obtained by concatenating the 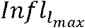 with its transpose. By cutting the resulting dendrogram in two, we obtain the two teams of the network.

#### Team strength, density and impurity

We calculate the team strength as follows:

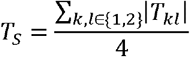

Where

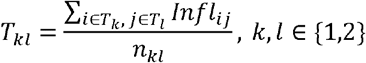

Density is calculated using the formula 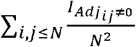, where *I*_*x*_ is the identity function that takes the value of 1 if x is true, 0 otherwise. Impurity is calculated using as the total fraction of inhibitory edges within teams and activating edges across.

### Simulating Networks using Boolean formalism

We simulated the TTNs primarily using a threshold-based Boolean formalism. Each gene in the network can take one of 2 states: +1 corresponding to high expression and -1 corresponding to low expression. We define a state vector S(t) to be a vector composed of the states of each gene part of the network at time ‘t’. If the network is composed of ‘N’ genes, then the state vectors have N elements, each representing the state of a gene. Following the update rule, the state is of the system updates via discrete transitions.

#### Update Rule

Adj is the Adjacency Matrix, in which the row indices are the Source Genes, and the column indices are the Target Genes. Adj_ij_ represents the interaction from the source gene ‘j’ to the target gene ‘i’

S(t) is the state vector at time ‘t’. S_i_(t) is the state of gene ‘i’

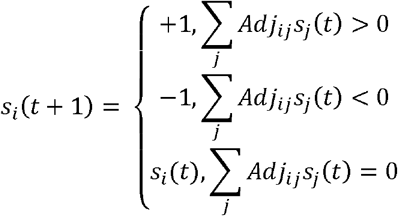

Where only one gene ‘i’ is randomly selected and updated at each time step.

We simulate the network starting at 100000 random state vectors. Each random state is updated using the dynamics given above until a steady state (i.e., S(t+1) = S(t)) is reached. The final state achieved from the simulation is then analysed.

Here, we follow the Asynchronous Update Rule, by which at each time step, a single gene is selected at random and its state is updated. To check whether a steady state has been reached, we used synchronous update.

#### ODE-based simulations – RACIPE

We performed the ODE-based simulations using a tool called RACIPE (Huang et al. 2018). Briefly, RACIPE constructs a system of ordinary differential equations based on the edge list of a network, where each equation describes the dynamics of a node *T*, activated by the set of nodes *P* and inhibited by the set of nodes in the network as follows:

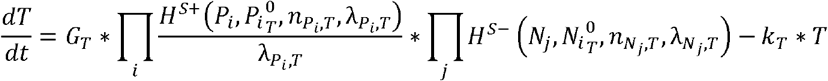

Here *T, P*_*i*_*and N*_*j*_ represent the concentrations at the given time.*G*_*T*_ and *K*_*T*_ denote the production and degradation rates, respectively 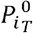 is the threshold value of *P*_*i*_.*n* is the hill’s coefficient and λ is the fold change in production rate due to a regulator. The regulation is modelled using a shifted hill function of the following form:

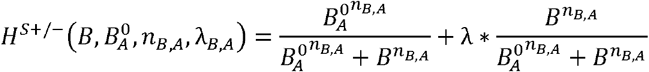

For this system of ODE models, RACIPE samples each individual parameter set uniformly from a pre-defined, biologically valid range, and numerically simulates them at multiple initial conditions until a steady state is reached. The final output consists of the collection of steady states for each parameter set. The steady states are then normalized and discretized as follows:

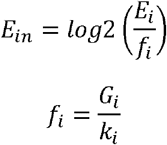

Where for the *i*^*th*^ node, *E*_*in*_ is the normalised expression level and *E*_*i*_ is a steady state expression levels. We then normalize the *E*_*in*_ values for each node across all steady states for all parameter sets, and discretize the positive values as 1 and negative values as 0.

### Metrics

#### F1 – Fraction of Pure States

A “pure” state is defined as a steady state in which genes belonging to one team have high expression and those belonging to the other team have low expression. F1 is defined as the ratio of total number of Pure States to the total number of steady states. In the case of embedded networks, the pure states were calculated for gene teams of the core.

#### CC AB – Correlation Coefficient between Average Expressions of Team A & B

The average expression of genes in Team A (Team A_Avg_) and the average expression of genes in Team B (Team B_Avg_) were calculated for each steady state. The CC AB was then calculated as the Spearman correlation coefficient was found for Team A_Avg_ vs Team B_Avg_.

### Generating artificial networks with different values of density, size and impurity

Artificial networks are generated using the stochastic block model (Holland et al. 1983), with constant probability of edges in all blocks of a network. The block model allows us assign team configurations. In TTN, the four blocks belong to two within-team interactions and two across-team interactions. We first generate four blocks of size NXN each, where N is the size of each team. Then, within each block, we add edges weighted according to the nature of the block (1 in within team interactions and -1 in across team interactions) with probability 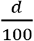 where d is the desired percentage density. We then change the weights of each edge with a probability 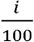, where “i” is the desired percentage impurity of the network. To investigate the effect of specific core and external network parameters, we generated Networks as follows.

#### Core Network

For Generating Core Networks with varying Edge Density, as per the density required, the number of Edges that need to be removed per quadrant of the core adjacency matrix (Adj) was decided as follows:

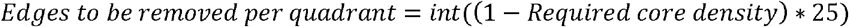

From each quadrant, the specific number was removed. So, the total number of edges in each quadrant is the same, i.e., the total number of incoming positive and negative interactions to one team is the same as the other. This was done to ensure that asymmetry in the number of edges doesn’t skew our initial experiments.

For the addition of impurity, networks of their respective densities are taken as input. Based on the required percentage edge impurity, the number of edges to be flipped from activation to inhibition and vice versa is decided as follows.

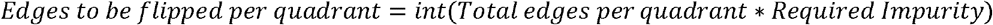

And in every quadrant, the same number of pure edges are flipped impure ones

#### Generating the External Network of a specific Mean Connectivity

Throughout our experiments, we have maintained a fixed size of 30 Genes for the external network. The total number of edges in the external network is determined based on the desired value of Mean Connectivity.

In the experiments to determine the effect of variation in the Mean Connectivity of the external Network, the Mean Connectivity was strictly controlled. In all other simulation experiments, the Mean Connectivity (m) parameter ranged between 2 and 12 for the external network.

Hence on average, every external node has **’m’** incoming/outgoing interactions. To achieve this, we first generate 30X30 matrix of zeroes, randomly select m^*^30 positions in the matrix and assigns 1 or -1 to these positions with equal probability, thereby ensuring an equal number of activations and inhibitions on average.

### Connecting the Core Network and the External Network

Now we have an external Network of 30 Genes and a two-team Core Network of 10 Genes (5 Genes Per Team). While connecting the External and Core Networks, the key parameters considered are the indegree and outdegree per Team.

#### Indegree

For a given value of indegree per gene in a team (i), it was ensured that, on average, each gene in each team had an indegree of (i). Furthermore, the indegree of each team was kept exactly the same. (Figure S8).

#### Outdegree

Similarly, for a given value of outdegree per gene in a team (o), it was ensured that, on average, each gene in each team had an average outdegree of (o). And the outdegree of each team was kept exactly the same.

#### Frustration

For a give state (*S*) of the network, an edge (*j* → *i*) is frustrated if *Adj*_*ij*_*S*_*i*_*S*_*j*_ < 0. The frustration of the state is then defined as the fraction of frustrated edges in the network corresponding to that state. In the core TTNs generated in this manuscript, the frustration of a pure state is zero in absence of any impurities, since nodes within the same team have the same sign, i.e., *S*_*i*_ * *S* _*j*_ >0 and the corresponding edge, if it exists, will be positive (*Adj*_*ij*_ ≥0). Similarly, for nodes belonging to different teams, *S* _*i*_ * *S* _*j*_ <0 *and Adj*_*ij*_ ≥0 for all (*i,j*) in absence of impurities. In presence of impurities, the only frustrated edges for pure states will be the impure edges, making the frustration of pure states equal to the fraction of impure edges in the network.

## Data and Code availability

The raw data files (network files, steady state frequency files) are available with the corresponding authors upon reasonable request. Processed data used to generate figures and codes to generate the networks and analyse the raw data has been deposited in the GitHub repository at https://github.com/saishyam1/Resilience_of_Teams.

## Author Contributions

SS and NN, VA, and KH performed analysis and data analysis, MKJ and KH conceptualized and supervised research. MKJ received funding. All authors contributed to writing and editing the manuscript.

