## Supplementary figures for "Mutually inhibiting teams of nodes: A predictive framework for structure–dynamics relationships in gene regulatory networks"

### Topological conditions of gene regulatory networks enabling robust binary cell-fate decision-making

Running title: Topological analysis of binary cell-fate networks

#### Abstract

Embryonic development entails multi-layer branching of cell-fate decision processes; often, each branch corresponds to a binary cell-fate decision. The two phenotypes at such branching points usually have mutually exclusive gene expression patterns, driven by two master regulators that mutually inhibit each other and can activate themselves, forming a toggle switch. The dynamics of a toggle switch have been extensively studied. However, the regulatory networks underlying cell-fate decisions often involve many transcription factors. We recently showed that these networks can be organized as two mutually inhibiting "teams of nodes", such that the nodes of the same team predominantly activate themselves, while those of the opposite teams primarily inhibit each other. While the ideal cases of fully connected and consistent networks of two "teams of nodes" have been well-understood, a detailed understanding of the effect of varying levels of connectivity and inconsistency in the edges concerning the configuration of teams is lacking. Here, we simulate networks of two teams (TTN) under varying degrees of edge density, edge impurity, and external influence to identify the conditions under which these networks can give rise to a robust binary cell-fate decision system. We find that the emergent cell-fate decision landscape is much more sensitive to inconsistent edges than to low connectivity, and that TTN generally leads to a more robust landscape than a toggle switch in the context of buffering external perturbations. We validate our results by simulating 42 GRNs from the cell collective database. Despite not all being cell-fate decision networks, coercing these networks to TTN format revealed strong correlations between the metrics of phenotypic landscape and edge impurity. Our results offer insights into the design principles of gene regulatory networks engaged in binary cell-fate decisions and can be extended to evaluate strategies for cellular reprogramming.

**Keywords**

Cell-fate decisions; Gene Regulatory Networks; Toggle Switch; Network analysis; Multistability

#### Supplementary Figures

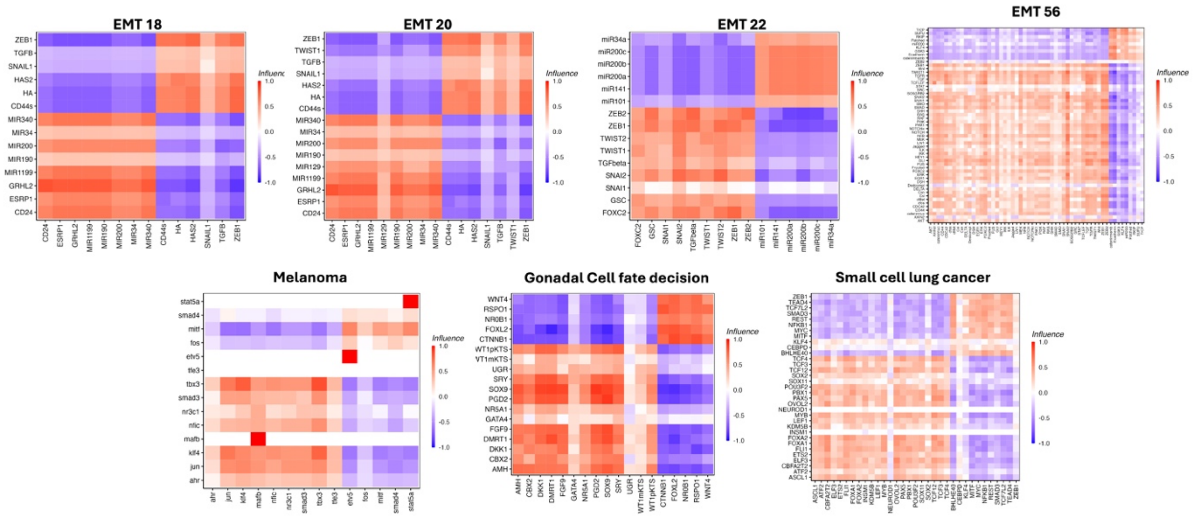


***Figure S1****: Influence matrices of cell fate decision networks underlying various contexts (mentioned in the title of each heatmap) used in Figure 1C.*

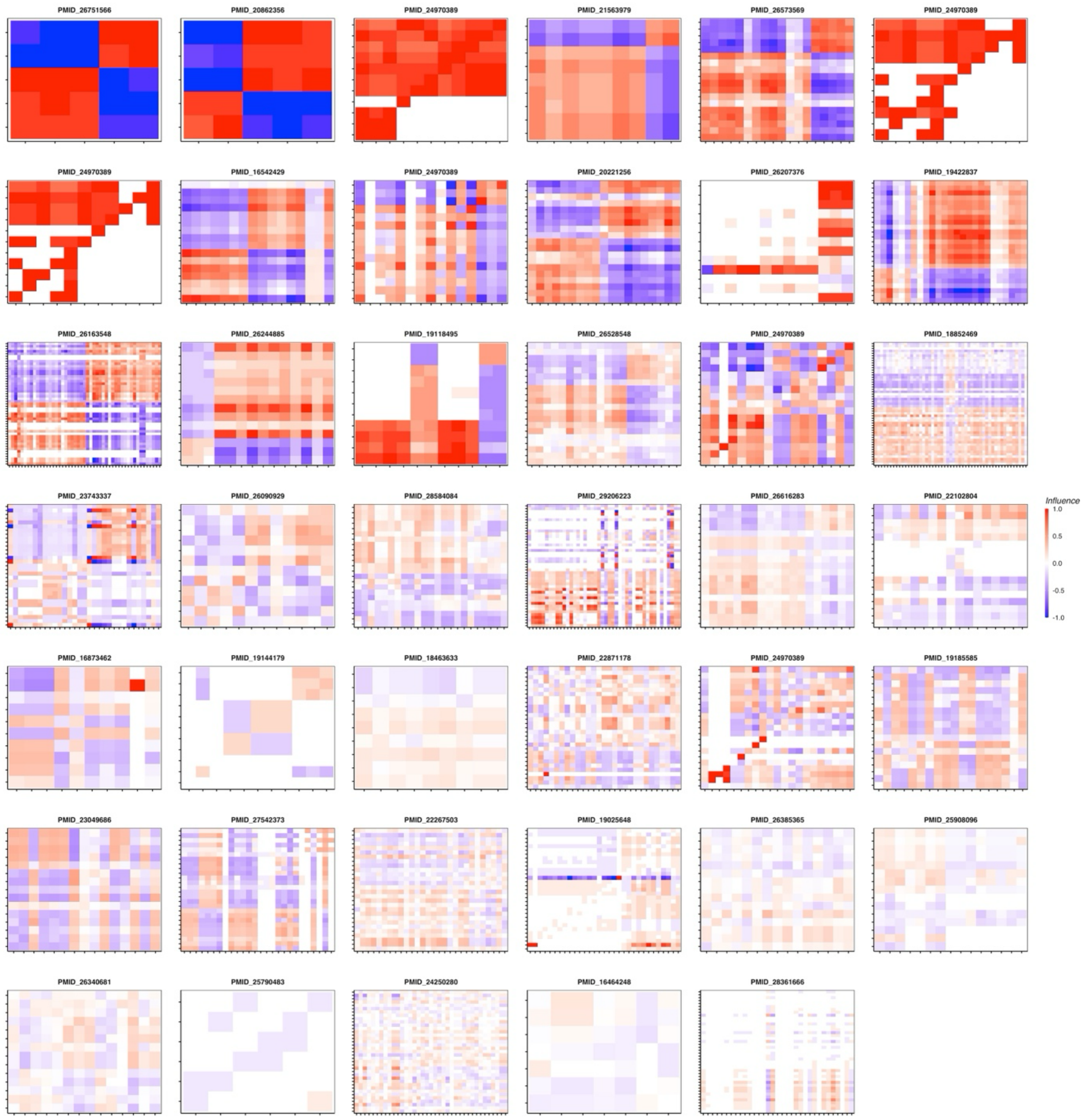


***Figure S2**: Influence matrices of the cell collective networks.*

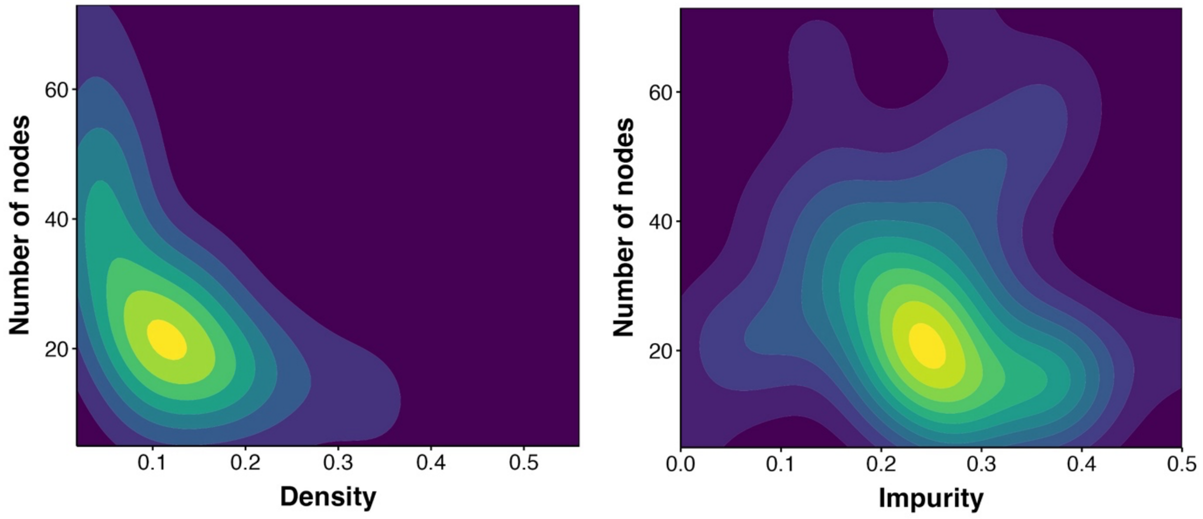


***Figure S3***: *Density maps showing the network size and edge density (left) and edge impurity (right) for cell collective networks*

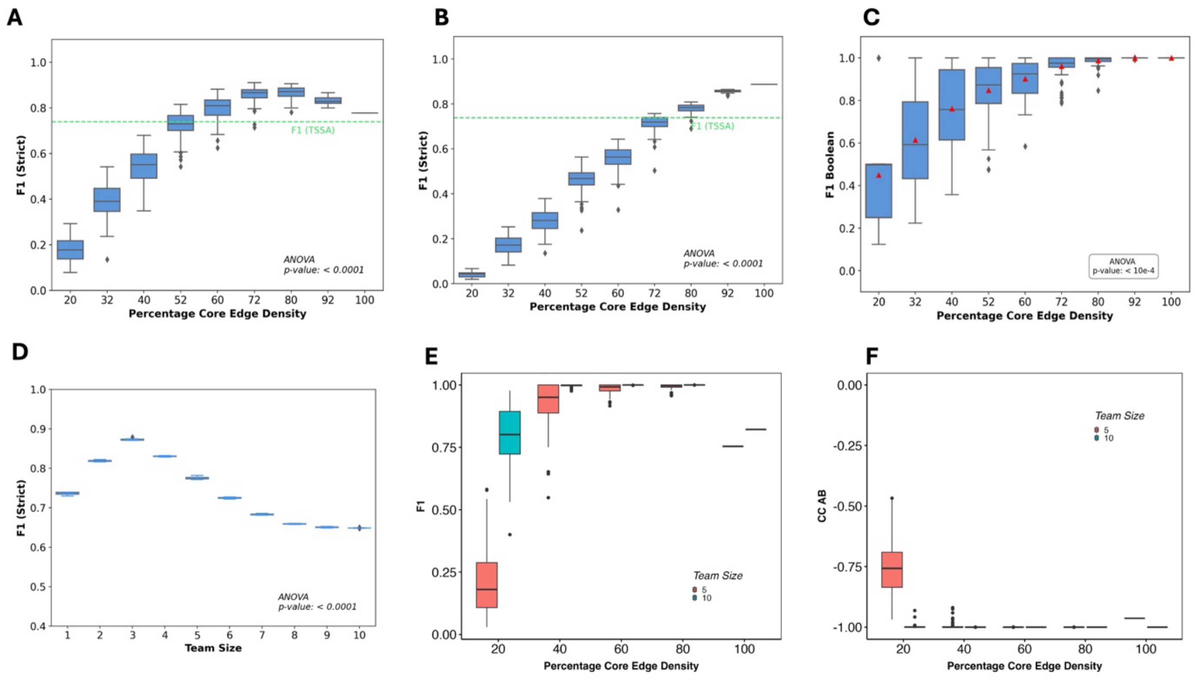


***Figure S4***: *Change in F1 and CC AB as a function of density and team size. A) Change in F1 calculated using RACIPE with change in TTN density. B) Same as A but for TTN without self-activation C) Same as B but for Boolean simulations D) Change in RACIPE F1 as a function of team size for 100% density TTN. E-F) F1 and CC AB with changing densities of TTN from Boolean simulations, for team sizes 5 and 10.*

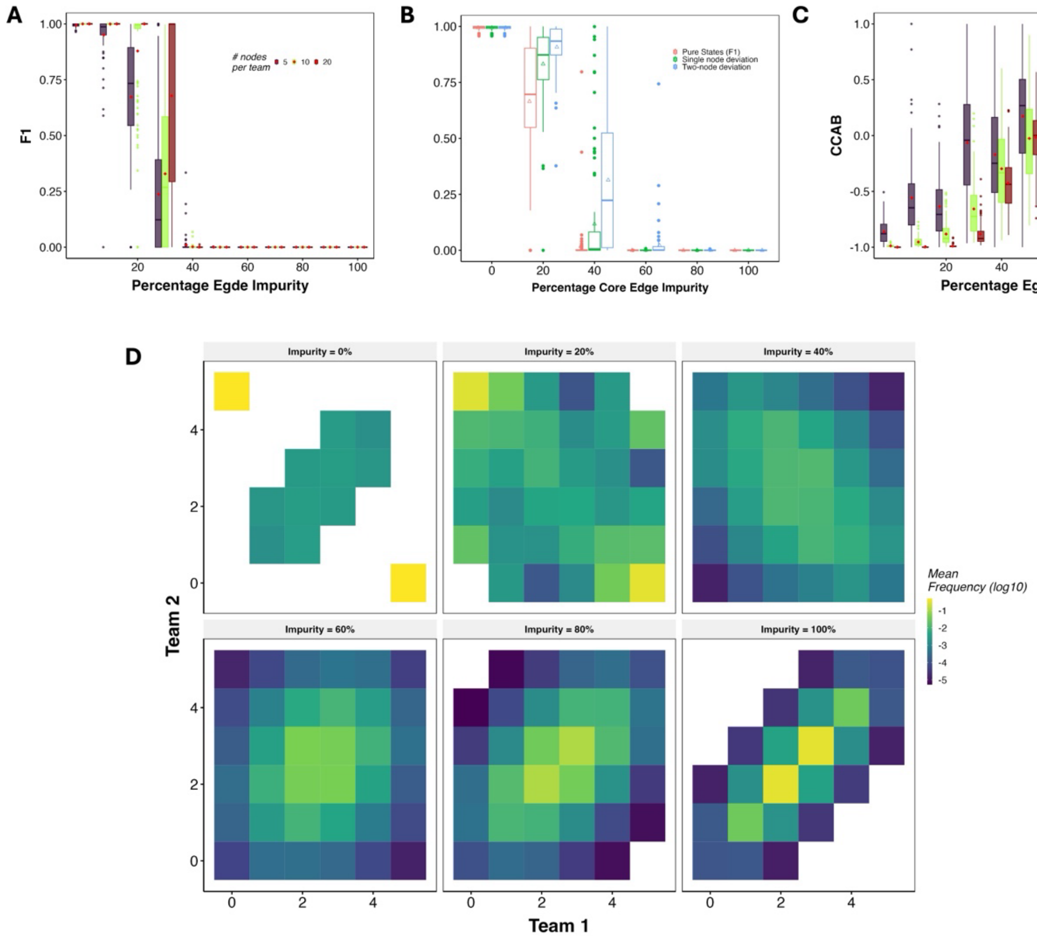


***Figure S5***: *Distributions of A) F1 and B) Relaxed F1 metrics and C) CC AB for TTS Core networks with different values for Percentage Core Edge Impurity at 80% Core Edge Density. D) Heatmap depicting the frequency landscape of Team A Score to Team B Score, for Networks of 80% Core Edge Density and different values percentage core edge impurity.*

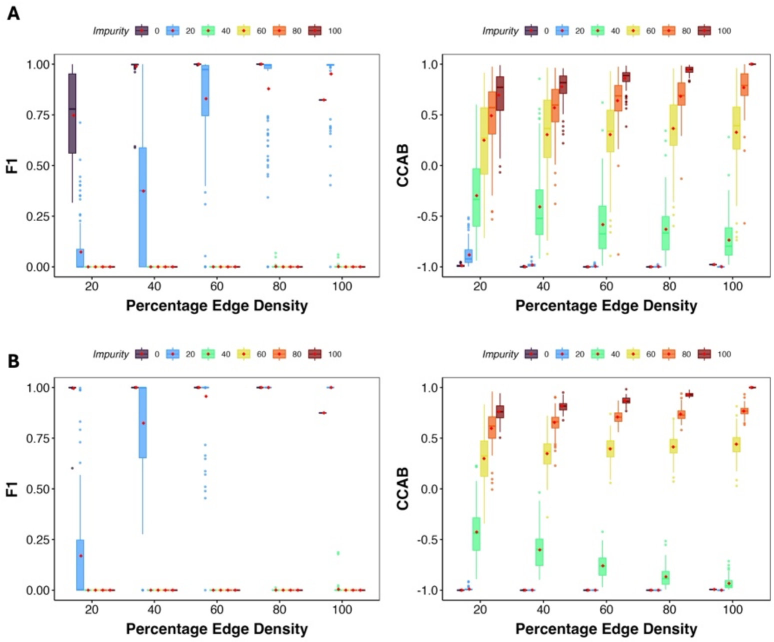

***Figure S6***: *Effect of Density and Impurity on F1 and CC AB of (A) 20 node and (B) 40 node TTN.*

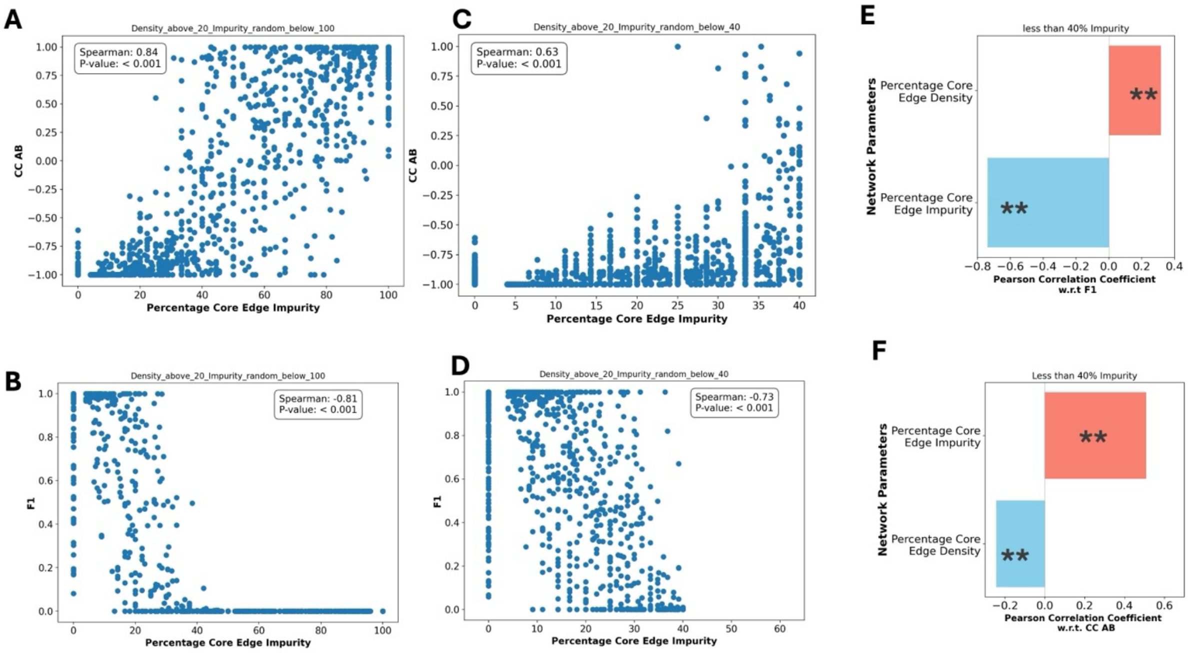


***Figure S7***: *Scatter plot of Percentage Core Edge Impurity vs A) CC AB and B) F1 of the corresponding network, for Networks with Core Edge Density greater than 20% and Impurity less than 40%. Scatter plot of Percentage Core Edge Impurity vs C) CC AB and D) F1 of the corresponding network, for Networks with Core Edge Density greater >= 20% and Impurity <= 100%. Scatter plot of log( Team Strength ) vs F1 of corresponding network for Networks with 80% Core Edge Density and Impurity <= 100%. Box Plots representing the Pearson's Correlation coefficient of Network Parameters, Percentage Core Edge Density, and Impurity with E) F1 and F) CC AB. ** indicates that p-value of t-test < 0.05.*

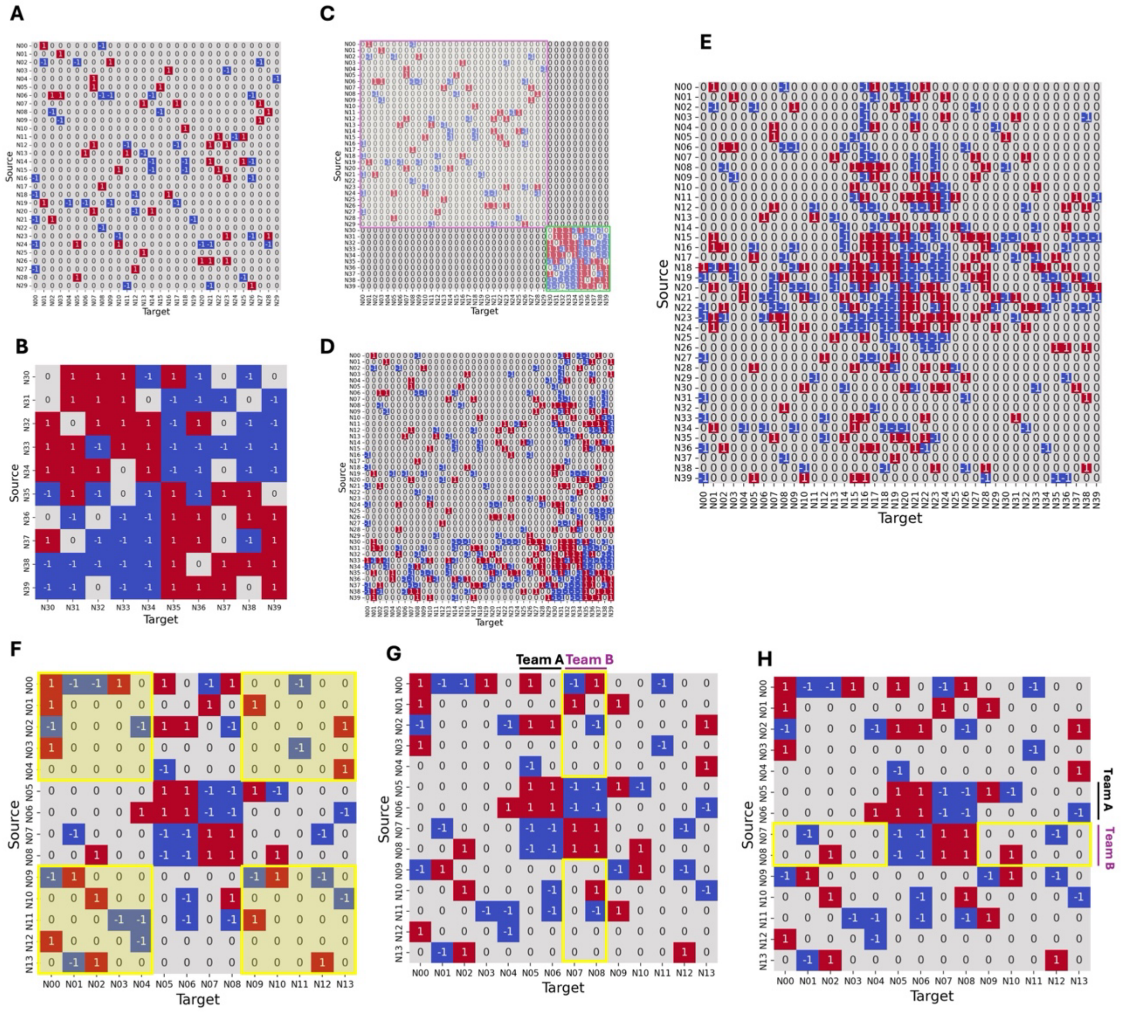


$$\textbf{Mean Connectivity of Peripheral Network} = \frac{\textit{Total Interactions Among Peripheral Genes}}{\textit{Number of Peripheral Genes}}$$

$$\textit{Total Number of Genes in Team B} = G_B$$

$$\textbf{Average Indegree per Gene}_{\textbf{Team B}} = \frac{\textit{Indegree of the Team B}}{G_B}$$

$$\textbf{Average Outdegree per Gene}_{\textbf{Team B}} = \frac{\textit{Outdegree of the Team B}}{G_B}$$

***Figure S8:*** *Schematic depicting the creation and key network parameters investigated in TTS embedded Networks. In the schematic, a TTS-2 is embedded in a 10-Gene External Network. A) Randomly generated Peripheral Network of 30 genes, N00 to N29, having desired Mean Connectivity (MC = 2.8 in A). B) Randomly generated Core Network of 80% Core Edge Density and 20% Density to be embedded, 10 Genes, N30 to N39. C) Bringing together the Core and Peripheral Networks. D) Introducing incoming and outgoing edges between the core and peripheral network. E) Centering the core shown in the adjacency matrix D by swapping the names of genes, N15 to N2, with genes N30 to N39. F) Yellow Boxes indicate the interactions between external genes. Mean Connectivity is calculated as the ratio of total interactions among external genes to the total number of external genes. In G), the Yellow Box highlights the incoming edges to Team B, i.e., the Indegree of Team B. The average indegree per gene of a Team is calculated as the ratio of the total number of incoming interactions towards genes in the team to the number of genes in the team. In H), the Yellow Box highlights the outgoing edges to Team B, i.e., the outdegree of Team B. The average indegree per gene of a Team is calculated as the ratio of the total number of outgoing interactions from genes in the team to the number of genes in the team.*

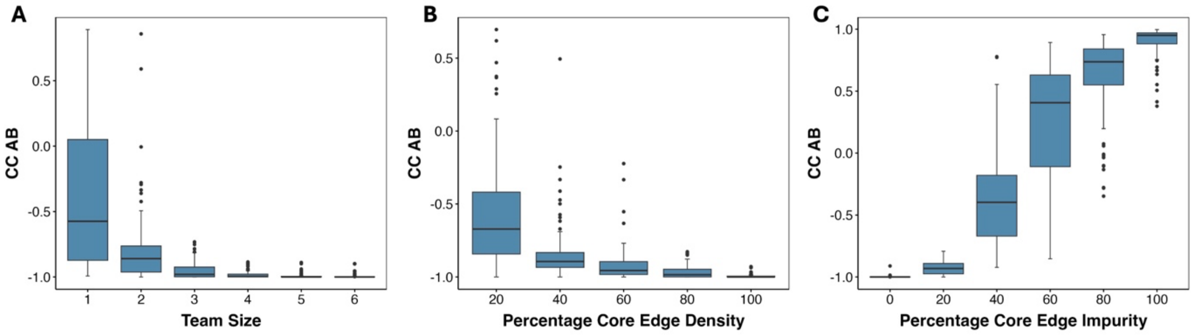


***Figure S9:*** *CCAB as a function of the A) Team Size, B) Density and C) Impurity for embedded TTNs.*

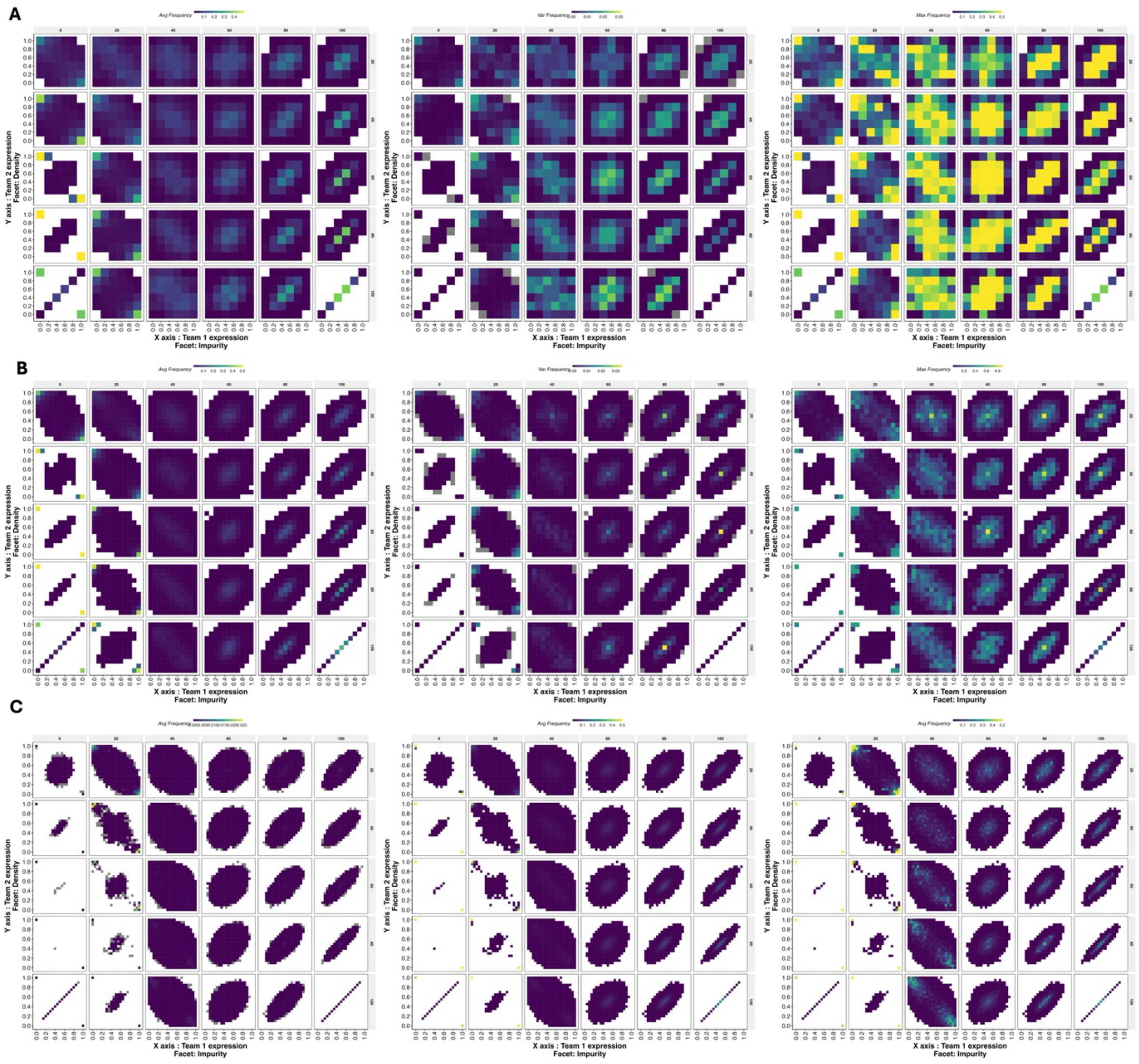

***Figure S10**: Heatmaps representing the abundance of different steady state compositions in the analyzed networks. There are 9 panels and each panel has 30 subpanels. Each row of the 3 X 3 figure corresponds to a team size: A) 5, B) 10 and C) 20. The left column corresponds to average frequency, middle column corresponds to*

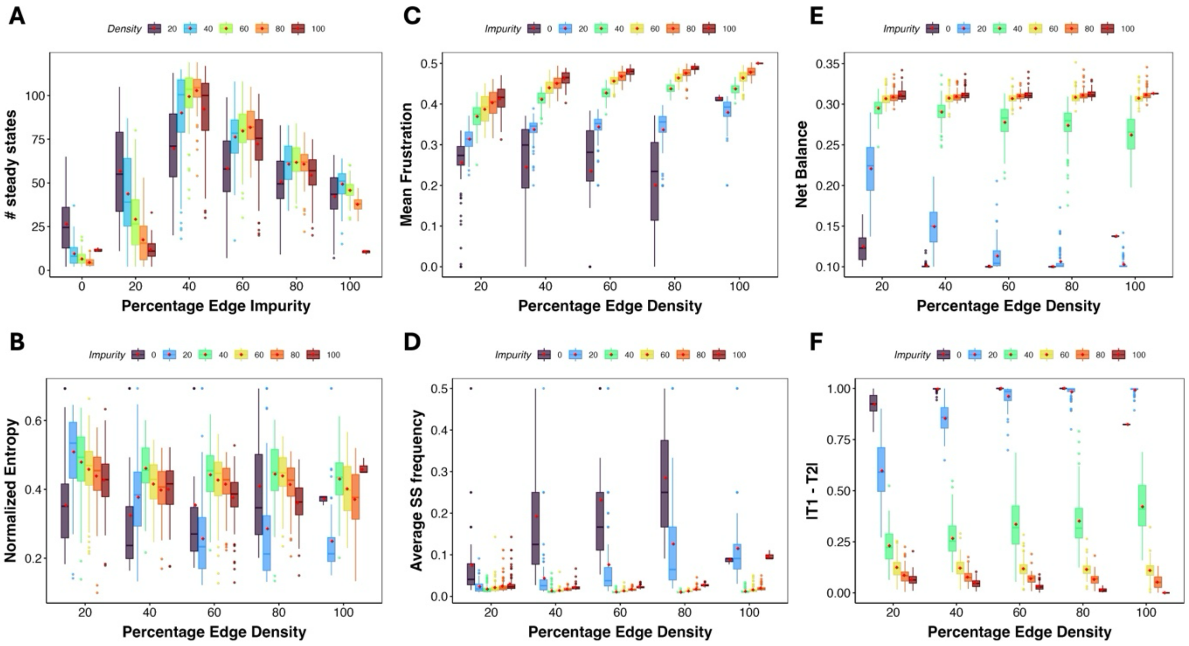


***Figure S11**: Effect of change in density and impurity on various measures of phenotypic landscape.*
